# The artemisinin-induced dormant stages of *Plasmodium falciparum* exhibit hallmarks of cellular senescence and drug resilience

**DOI:** 10.1101/2023.01.29.526019

**Authors:** Jaishree Tripathi, Michal Stoklasa, Sourav Nayak, Kay En Low, Erica Qian Hui Lee, Laurent Rénia, Benoit Malleret, Zbynek Bozdech

**Affiliations:** School of Biological Sciences, Nanyang Technological University, Singapore, 637551; Department of Microbiology and Immunology, Yong Loo Lin School of Medicine, National University of Singapore (NUS), Singapore, 117597; Electron Microscopy Unit, Yong Loo Lin School of Medicine, National University of Singapore (NUS), Singapore, 117597; Lee Kong Chian School of Medicine, Nanyang Technological University, Singapore, 636921; A*STAR Infectious Diseases Labs (A*STAR ID Labs), Agency for Science, Technology and Research (A*STAR), Singapore; Singapore Immunology Network (SIgN), Agency for Science, Technology and Research (A*STAR), Singapore, 138648

## Abstract

Recrudescent infections with human malaria parasite, *Plasmodium falciparum*, presented traditionally the major setback of artemisinin-based monotherapies. Although introduction of artemisinin combination therapies (ACT) largely solved the problem, the ability of artemisinin to induce dormant parasites still poses major obstacle for current as well as future malaria chemotherapeutics. Here, we developed a robust laboratory model for induction of dormant *P. falciparum* parasites and characterized their transcriptome, drug sensitivity profile and cellular ultrastructure. We show that *P. falciparum* dormancy requires a ~5-days maturation process during which the genome-wide gene expression pattern gradually transitions from the ring-like state to a highly unique form. The transcriptome of the mature dormant stage carries hallmarks of cellular senescence with downregulation of most cellular functions associated with growth and development, but upregulation of selected metabolic functions and DNA repair. Moreover, the *P. falciparum* dormant stage is considerably more resistant to essentially all antimalaria drugs compared to the fast-growing asexual stages. Finally, the unique cellular ultrastructure further suggests unique properties of this new developmental stage of the *P. falciparum* life cycle that should be taken into consideration by new malaria control strategies.

## Introduction

Artemisinin combination therapy (ACT) is currently the backbone of all malaria elimination programs around the world^1^. Since their implementations in the early 2000’s, ACTs have contributed considerably to the reduction of malaria morbidity and mortality achieved in the first two decades of the 21^st^ century worldwide. Unfortunately, the decreasing efficacy of ACT emerged in eastern part of the Greater Mekong Subregion as early as 2009^2,3^ and now emerging in other parts of the world including sub-Saharan Africa^4–6^ is posing a major concern for the future^7,8^. Better understanding the ACT mode(s) of action as well as their resistance mechanisms is still urgently needed not only for managing their current clinical implementations but also for rational designs of new strategies^9^.

All currently used ACTs are composed of two main components: (i.) fast acting artemisinin or its derivatives (further artemisinins), mediating the removal of majority of the parasites during the early stages of the treatment course; (ii.) the long lasting/acting partner drugs that eliminate the residual parasite loads remaining in the patient during the later stages of the treatment^10^. The rationale for this ACT design is based on the short bioavailability of artemisinin that despite its high antiparasitic potency, often fails to eliminate all parasite loads^11,12^. Indeed, in earlier clinical monotherapy-based implementations, artemisinins were shown to be highly effective in rapid reductions of parasite loads in uncomplicated *P. falciparum* infections yielding complete parasite clearance after 5 to 7-days treatment period (reviewed in^13^). Despite this, considerable recrudescence was reported essentially in all the clinical studies with wide range of potency (3-50%)^14^, depending on geographical and epidemiological background. Interestingly, the rate of recrudescence appears to be linked predominantly with the overall parasite load at the start of the treatment rather than other epidemiological or clinical factors^15^. This indicated that the recrudescing infections rise from (some form of) a residual parasite fraction that remains in the patients’ body after the 5-7 day treatment. It is plausible that this “residual” parasite population consists of unique parasite forms that are resilient to further artemisinin treatment and can persist in the patients’ body for extended periods of time. The most fitting hypothesis for such parasite form is a quiescent (or senescence-like, see below) state that was previously observed in other microbial pathogens such as bacteria^16^.

Indeed, exposure of *P. falciparum* to artemisinins *in vitro* induces dormancy-like forms that could represent this residual parasite (sub)population^17^. Initially reported by Kyle and Webster in 1996, treatment of the ring stage *P. falciparum* parasites with clinically relevant concentrations of artemisinins induced dormant parasites that can be detected as pyknotic-like cellular forms in an *in vitro* culture for 3-8 days^18^. Exploring this further, the same group established a tractable laboratory model in which treatment of ring stages with 700nM (~200ng/ml) of dihydroartemisinin (DHA) for 6 hours (hr) drives parasites to dormancy that persists in an *in vitro* culture for up to 20 days^19^. In the following series of studies, it was shown that varying levels of drug resistance can affect the rate of the dormancy-dependent recrudescence^20^. The persister parasites exhibit an altered pattern of development^21^, expressing several genes related to metabolism^22^ and cell cycle regulation^23^. Moreover, the dormant parasites are characterized by condensed cytoplasm and nucleus but enlarged mitochondrion that retains its membrane potential^24,25^. Indeed, there is a distinct rearrangement of the dormant parasite cellular structure with a distinct, new mitochondrial-nuclear associations^25^. It was suggested that restructuring is likely induced by an oxidative stress and might lead to an altered transcriptional activity within the nucleus. In addition to the *in vitro* observation, there is now mounting evidence that the persister parasites also occur *in vivo*. Initially, dormant forms were shown to be induced by artemisinin in the *in vivo* rodent malaria model *P. vinckei*^26^ but were also clearly detected in the blood of volunteers in control human infection studies infected with wild type and PfK13 mutant/artemisinin resistant *P. falciparum* strains^27^.

Taken together, these results suggest that persister forms of *P. falciparum* parasites could provide the reservoir for recrudescent infection after artemisinin monotherapy and as such play a role in the current ACT failures (*see above*). Here, we developed a robust culturing system for artemisinin-induced dormant parasites (of *P. falciparum*) that allowed us to analyze their basic biological blueprint in greater detail. Using transcriptomic analyses, we showed that the induced dormant stage of *P. falciparum* represents a unique physiological state requiring 4-5 days of maturation. The *day 5 mature dormant persistors* (d5MDP) exhibit highly specific and unique transcriptome, but also highly irregular ultrastructure as well as drug sensitivity profiles. Cellular and molecular features of the d5MDPs are consistent with their role as a residual parasite population responsible for recrudescence.

## Results

### Cellular definition of the artemisinin-induced *P. falciparum* dormancy developmental cycle

To gain further insights into the biological processes that drive induction and persistence of the dormant forms of *P. falciparum* parasites and the subsequent re-emergence of regular IDC stages, we adopted the culturing protocol originally developed by Teuscher, F. *et. al*. 2010 for induction of *P. falciparum* dormancy^19^ with few marked modifications. Briefly, synchronized *P. falciparum* culture (3D7 strain) was treated at the ring stage (6-12 hours post invasion, HPI) by 700 nM dihydroartemisinin (DHA) and subsequently cultured for 12 days. During the first three days, magnetic-assisted cell sorting (MACS) was used to remove mature stages of the parasite intraerythrocytic developmental cycle (IDC), trophozoites and schizonts (**Figure 1A**). Each day, the distribution of morphological stages and parasitemia was evaluated by microscopy (Giemsa-stained blood smear) and Fluorescence-Assisted Cell Sorting (FACS) (**Figure 1B-1E; Figure S1**,**Material and Methods** section).

**Fig 1.**
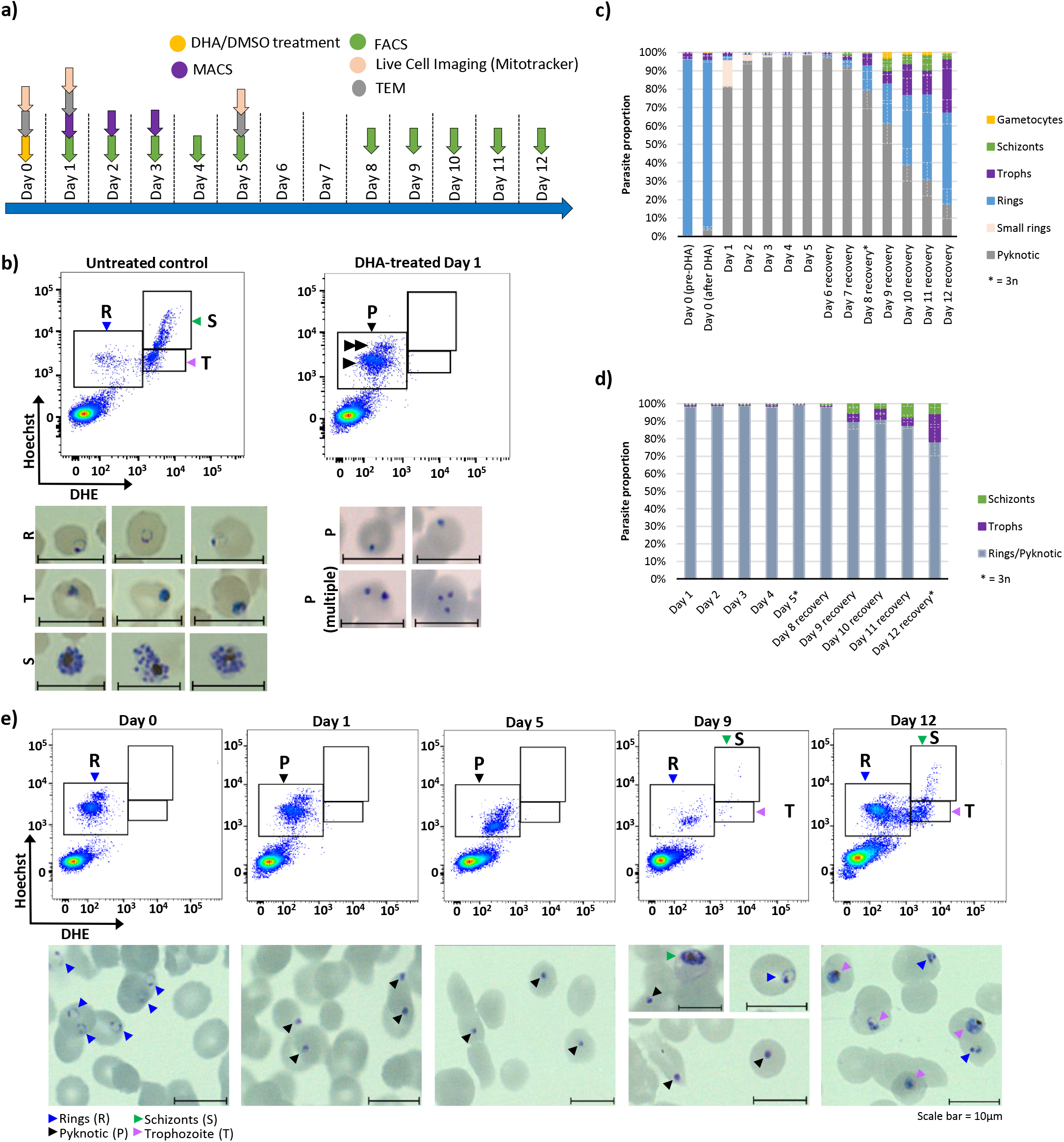
Dihydroartemisinin (DHA) Induced Dormant Stages of *P. falciparum* Parasites. **a)** A schematic diagram showing the experimental design. Briefly, 500μl of packed RBCs infected with *P. falciparum* (2-8% parasitemia, 6-12 HPI) were treated with 700nM DHA for 6 hours to induce pyknotic dormant forms (Day 0). Possibly resistant parasites were removed from the culture by passing the culture through MACS columns on three consecutive days (Day 1-3). Fresh blood was added on Day 5 for supplementation and prevention of lysis of RBCs. Thereafter, 500-cells were sorted from the dormancy-induced culture for transcriptomics analysis by FACS on Days 1-5 and Days 8-12. Samples for live-cell imaging and TEM were collected on Day 0, Day 1 and Day 5. **b)** A flow cytometry dot plot showing the gating strategy and sorting of parasites via FACS. An untreated control culture (left panel) with mixed IDC stages is shown in comparison to the DHA-treated culture on Day 1 (right panel). For distinguishing various *P. falciparum* stages, Hoechst 33342 and DHE staining was performed. To confirm the correct gating of different morphological stages, 3000-5000 infected RBCs were sorted into RPMI media and diluted into 0.3-0.5% parasitemia for the Giemsa smears shown here (see **Materials and Methods**). R = rings, T = trophozoite, S = schizonts, P = pyknotic, single arrow = infected RBCs with single pyknotic parasites, double arrow = infected RBCs with multiple pyknotic parasites. Scale bar = 10μm. Proportion of different parasitic stages evaluated by: **c)** Microscopy of Giemsa stained blood smears where 300 cells were counted for each day in four independent biological replicates. Error bars represent SEM. **d)** FACS, where 50,000 events were collected each day in four independent biological replicates. Error bars represent SEM. Asterisk (*) in c) and d) = only three biological replicates used. **1e)** Gating and sorting strategy for monitoring of the parasite dormant stages induction and recovery process. The FACS plots shown represent DHE + Hoechst 33342 staining of synchronized parasite culture after excluding debris and doublets. Giemsa smears of the representative parasitic stages in culture on corresponding days are shown below the FACS plots with colored arrows pointing to the different parasite stages. Scale bar = 10μm.

Using the Giemsa-stained blood smears we observed that one day after DHA treatment (Day 1), 81.8 % of the parasites (average of four bioreplicates, **Fig 1C**) exhibited a pyknotic morphology, that is characterized by densely stained, small, round-shaped cell-like structures (**Figure 1B, right panel, and Figure 1E, lower panel**). These changes contrast the (initial) ring stage parasites that are characterized by the “classical” O-ring shape with a single dense formation along the periphery, representing the parasite nucleus (**Figure 1B, left panel, and Figure 1E, lower panel**). The morphological transformation from ring-shape to pyknotic cells after exposure to artemisinin drugs is analogous to previous studies which have reported formation of rounded morphology with blue-stained cytoplasm and red-stained chromatin in Giemsa smears of dormant parasites^19,20^. Importantly, the proportion of the pyknotic parasites rose from 95.8 % to 98.9 % (in four bioreplicates) of the total parasitemia between Day 2 and Day 5; essentially dominating the entire parasite population within the culture (**Figure 1C**). The proportion of pyknotic forms remained >92% until Day 7, after which we could detect a rising number of asexual IDC stages (**Figure 1C**). This indicates re-emergence of the asexual growth that resumed fully by Day 10, with 60.2 % (average of four bioreplicates) of the parasites in one of the asexual IDC forms (**Figure 1C**). Crucially, this temporal morphological distribution was reproduced four times, each time with highly similar counts of the individual parasite morphologies on each day (**Figure 1C**, error bars). To complement the Giemsa stained blood smears, we also carried out FACS-based analysis of the parasite cultures using Hoechst-Dihydroethidium (Hoechst-DHE) double-staining protocol^28^, that can distinguish between the key IDC stages (rings, trophozoites and schizonts) (**Figure 1B and 1D and Figure S1B**). In this Hoechst-DHE-FACS methodology, the pyknotic and ring stage parasites exhibit an identical fluorescence intensity staining and hence fall within the same gate (**Figure 1B and 1E, Figure S1C** (FACS sorted smears)). Upon comparison, the parasite staging by FACS (**Figure 1D**) closely mirrored the parasite counting performed manually by Giemsa staining (**Figure 1C**) with >90% of the parasites (average of four bioreplicates) falling in the pyknotic-rings gate from day 1 to day 8 and recovery observed from day 9 onwards (**Figure 1D**). The purity of each parasite stage isolated from various FACS gates was confirmed by Giemsa staining of blood smears made from the FACS sorted fraction of the cells (**Figure S1C**). At this point, it is unclear whether the pyknotic parasite forms persisting in the cultures (~17 %) at Day 12 (**Fig 1C**) reflect longer persistent cells or dead parasites.

It is also important to note that during the first 4-5 days of the DHA-induction experimental time-course, the total parasitemia remained constant, staying at the initial level prior to the DHA treatment (**Figure S1A).** The subsequent “dips” in parasitemia reflect the necessary supplementations of the culture with fresh red blood cells (RBCs) on day 5 to counteract hemolysis. This suggests that the 6 hr treatment of the *P. falciparum* rings at 6-12 HPI with 700 nM of DHA does not lead to parasite elimination but instead to a uniform transition to pyknosis. This contrasts the treatment of 6-12 HPI rings with chloroquine (CQ, 250nM (10xIC_50_) for 24 hours) where a progressive reduction in parasitemia was observed from Day 0 (treatment) to Day 3 (**Figure S2A**). Mixed parasite morphology with both pyknotic forms and irregular/blebbed forms were observed as early as 24 hours after CQ treatment with progressive parasite elimination from the culture by Day 3 (**Figure S2B and S2C**).

### Transcriptome profiling of *P. falciparum* pyknosis uncovers day 5 mature dormant persistors (d5MDP)

Next, we took the advantage of the developed culturing systems and reconstructed the global transcriptome of the pyknotic cells along the entire 12-day time course. The main motivation was to evaluate if these pyknotic cell-like forms represent the genuine dormant stage(s) of *P. falciparum* and if so, what is their biochemical and cellular physiology. For this, we used the above-mentioned FACS-based cell isolation strategy coupled with “few-cells” RNA sequencing that allows morphology/stage-associated transcriptomic analysis of a defined number of *P. falciparum* cells (including single cell)^29^. Briefly, 500-parasite cells were captured from the pyknotic/ring stage FACS gate each day and subjected to SMARTseq2-based RNA sequencing for whole transcriptome profiling^30^ (for more details see **Material and Methods**). Due to the need of RBC supplementation on Day 5 (**Figure S1A**), the transcriptomic time course experiments were conducted in three separate phases: *induction and maturation* (further only *maturation*) between Day 0 to Day 5; *persistence* between Day 6-7; *re-emergence* of asexual growth between Day 8-12. While the transcriptome analyses of the *induction* and *re-emergence* phases were carried out in a single time course, the *persistence* phase was conducted in a separate experiment.

In the *maturation* and *re-emergence* transcriptomic dataset, we detected between 2152 to 3086 genes expressed in at least 80% of the timepoints representing the gene expression profile in at least one of the three biological replicas. Subsequently, we applied Fourier transformation to the identified gene expression profiles to generate an overview of the global transcriptomic pattern of the presumed *maturation* and *re-emergence* phases of *P. falciparum* dormancy. Using one of the biological replicates (i.e. “BR_C”) as the representative dataset, we identified at least 786 genes that exhibited gradual downregulation in their mRNA levels during the *maturation* phase reaching their minimum expression between Day 3 to 5 (**Figure 2A**, *green bar*). Crucially, most of these genes exhibited (re)induction from Day 8 onwards reaching its original ring-specific levels by Day 12. On the other hand, 1218 genes exhibited a progressive increase in expression from Day 1 to Day 5, with their peak expression exceeding their initial (Day 0 i.e. rings) mRNA levels starting Day 3. Analogously, mRNA abundance of these genes fell back to the initial (low) levels on Day 10 through 12 (**Figure 2A**, *red bar*). These results illustrate a dynamic expression pattern of more than 2000 (~35%) *P. falciparum* genes throughout the 12-day time course after the DHA-induction of pyknosis. This pattern is particularly surprising during the *maturation* phase (Day1-5) where we observed only gradual transcriptional changes contrasting the morphological profile with essentially all parasites in the pyknotic state (**Figure 1C to 1E**). This suggests that during the first 5 days, the DHA-induced pyknotic forms of *P. falciparum* undergo a developmental process where the initial ring-like transcriptional profile gradually transitions into a unique pattern (see below). Curiously, while the Day 1 (and to some degree Day 2) parasites relate transcriptionally to the ring stage, the transcriptional profile of the dormant/pyknotic parasites from Day 3 through Day 8, exhibit no resemblance to any stage of the asexual and sexual growth (**Figure 2C**). Crucially, this correlation pattern is well preserved in all three biological replicas of the *maturation* and *re-emergence* phases (**Figure S3A and S3B**), This also applies to the transcriptome of the *persistence* phase between Day 5-7 that bears no resemblance to any IDC or gametocytogenesis bound developmental stages (**Figure 3A & 3C and Figure S3C**). Moreover, there is a remarkably high correlation between the corresponding transcriptomes across the biological replicas across all three phases of the dormancy development, suggesting the specificity and reproducibility of the dormant parasite transcriptional characteristics (**Figure S4** and **Figure 3C**). This suggests that the observed transcriptomic reshuffling may represent an orchestrated transcriptional program that is “hardwired” into the *P. falciparum* genome, giving rise to a new previously undefined developmental stage. The concept of gradual transition might be also partly true for the *re-emergence* phase (Day 9-12), however, the 500-cell samples collected during this phase contained a progressively increasing number of morphologically distinct rings (**Figure 1C to 1E**). Taken together these data demonstrate that the DHA-induced dormant stages of *P. falciparum* represent a unique physiological state with transcriptional rearrangement of a broad spectrum of cellular and metabolic pathways (**Figure 2D & 2E**, *see beloW*).

**Fig 2.**
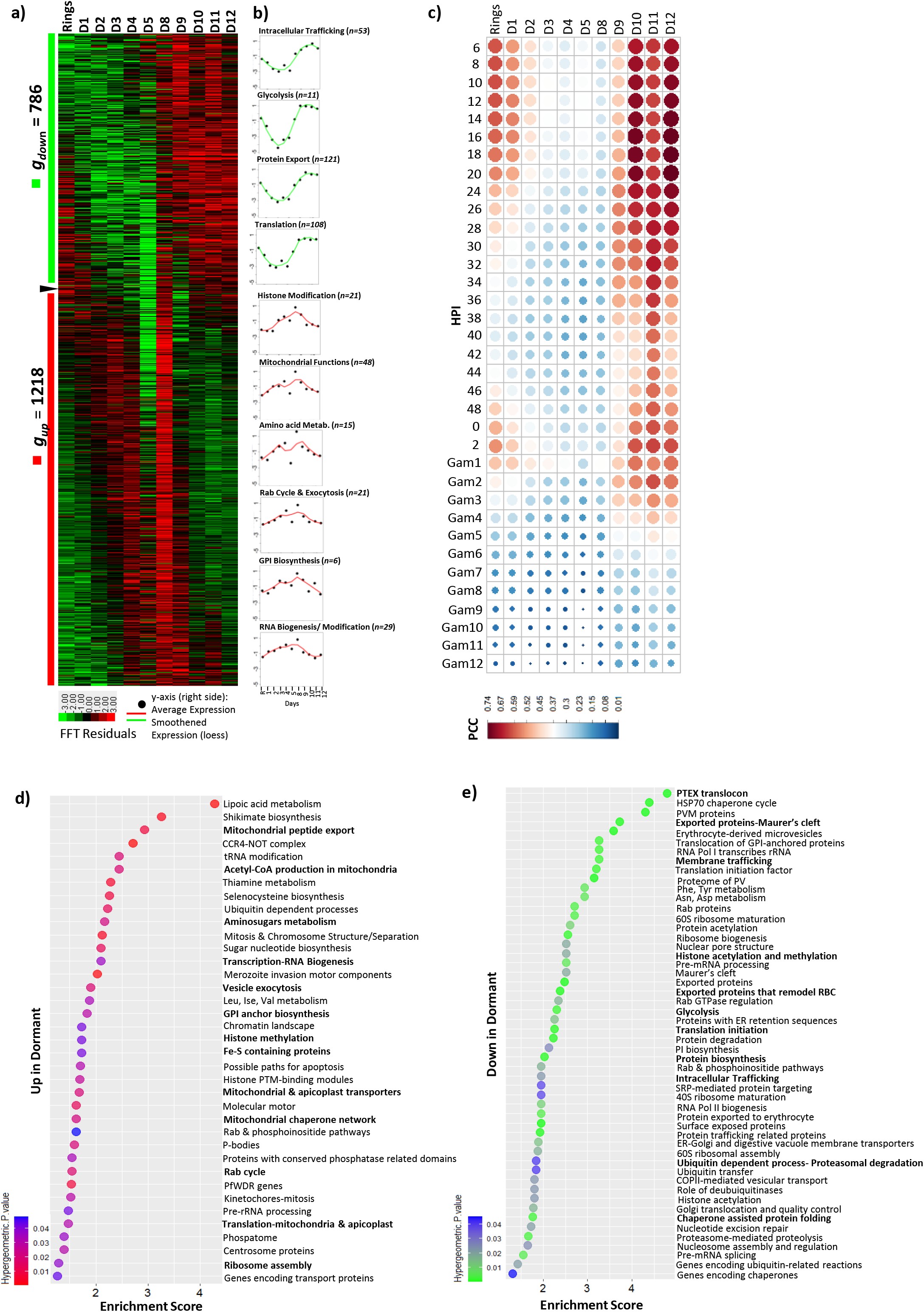
Transcriptomic profiling of dormancy developmental cycle in DHA treated *P. falciparum*. **a)** A heatmap depicting the Fourier transformed gene expression profile of *P. falciparum* during the *induction* and *re-emergence* phase of DHA induced dormancy. Genes upregulated and downregulated during Day 1 to 5 are highlighted by red bar (***g_up_***) and green bar (***g_down_***) respectively. **b)** A line graph showing the average expression (smoothened) of key cellular pathways enriched amongst the upregulated (red) and downregulated (green) genes between Day 1 to 5. **c)** A bubble plot depicting Pearson correlation coefficients (PCC) of *P. falciparum* transcriptomes obtained during the *induction-maturation* (Day 1 to 5) and *re-emergence* (Day 8 to 12) phases of dormancy in comparison to the high-resolution 3D7 reference transcriptomes for asexual (0-48 HPI) and sexual stages. **d)** and **e)** Bubble plots depicting pathways enriched amongst genes upregulated (d) and downregulated (e) during Day 1 to 5 after DHA induced pyknosis. All pathways are ordered by enrichment score and the color (red and green) denotes the statistical significance (hypergeometric p-value).

**Fig 3.**
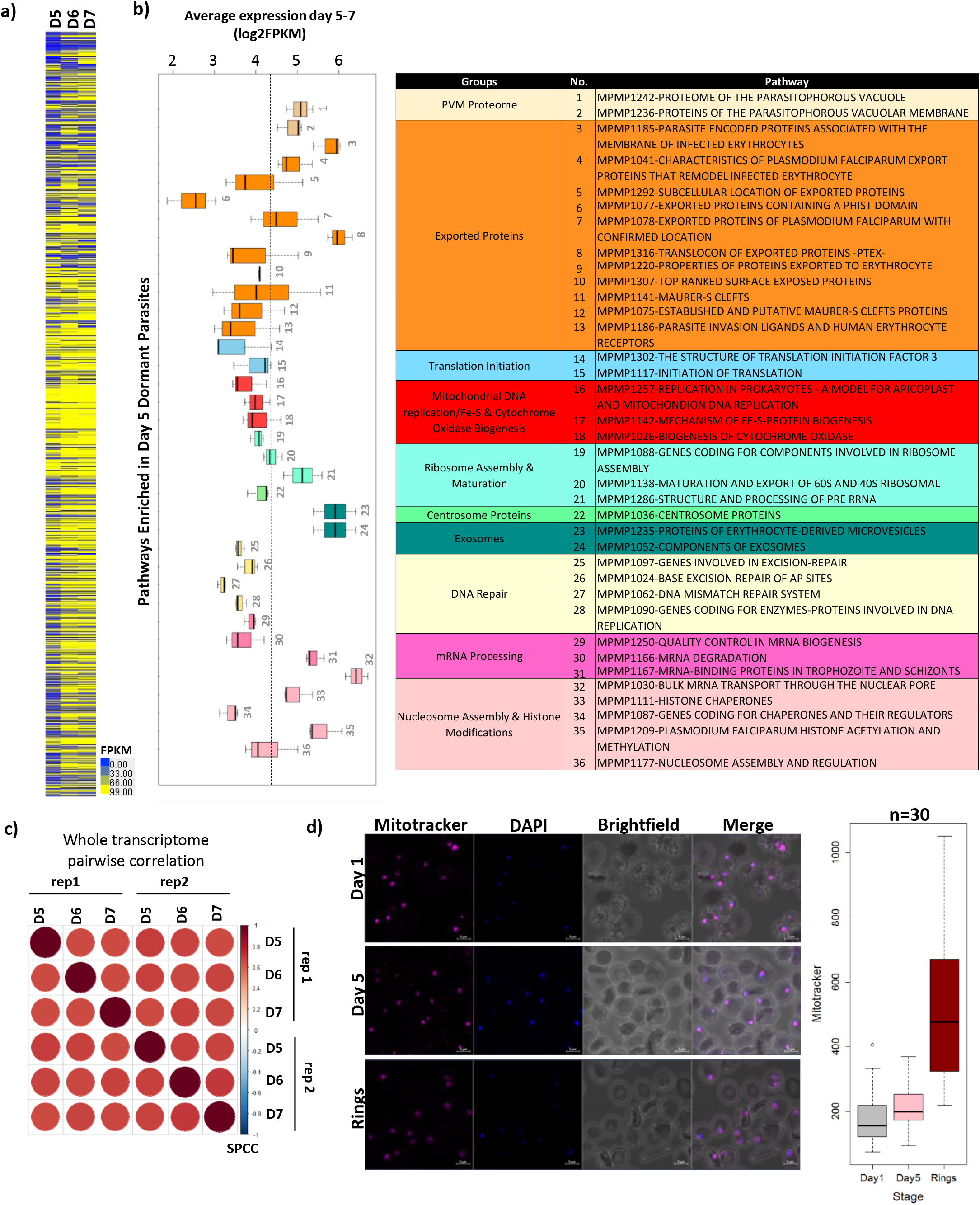
Transcriptome and Functional Pathway Enrichment for day 5 dormant parasites (d5MDPs). **a)** A heatmap showing expression for genes (***g_down_*** and ***g_up_*** in Fig 2a, same order) during the *persistence* phase (Day 6-7). Yellow color depicts higher FPKM values whereas blue depicts lower FPKM. **b)** A boxplot depicting average expression (in log2FPKM) of pathways found to be significantly enriched during Gene Set Enrichment Analysis (GSEA) of d5MDP transcriptome. The boxplots are color coded and categorized according to cellular/metabolic function similarities within the parasite (table on right). **c)** A bubble plot depicting whole transcriptome correlations (Spearman’s Rank Correlation Coefficients, SPCC) for Day 5,6 and 7 *persistence* phase of the parasite transcriptome (biological replicates - rep 1 and rep 2). **d)** Live-cell imaging of mitochondrial activity of dormant parasites. Untreated rings are compared with DHA-treated pyknotic parasites on Day 1 and Day 5 through mitochondria staining with Mitotracker Deep Red^Tm^ dye and nuclear staining with DAPI. The boxplot shows average intensity of Mitotracker Deep Red^Tm^ stain quantified in Day 1, Day 5 and ring stage parasites (n=30) using Zeiss ZEN software.

Based in these transcriptomics results, it appears that the dormant stage appears to require 4-5 days for a maturation process represented by the gradual transcriptional transformations. For that reason we wish to term the mature dormant stages the five (<u>5</u>)<u>d</u>ays <u>M</u>ature <u>D</u>ormant <u>P</u>arasite stage (5dMDP). The proposed concept of transcriptional retuning of the 5dMDP is also supported by functional enrichment analyses that revealed their specific gene expression pattern with inductions/suppressions of several key metabolic and cellular functions. First, we observed a strong upregulation of a broad spectrum of genes involved in mitochondria-linked energy metabolism including ATP synthase subunits, cation-transporting ATPase 1, citrate/oxoglutarate carrier protein, cytosolic Fe-S cluster assembly factors and several mitochondrial and apicoplast ribosomal proteins during day 1 to day 5 (**Figure 2B**). Additionally, several pathways linked to RNA biogenesis, GPI biosynthesis, histone modifications, mitosis and chromosome separation showed increased average expression during the dormancy *maturation* phase (**Figure 2B, lower panel**). These are complemented by an array of specific cellular pathways such as mitochondrial peptide export, Acetyl-CoA production in mitochondria, amino-sugar metabolism, mitochondrial & apicoplast transporters, etc., all of which were found to be enriched amongst the genes induced during the 5dMDP maturation (**Figure 2D**). Inversely, glycolysis but also factors of intracellular trafficking, protein export and translation were found to be suppressed through the 5dMDP maturation (**Figure 2B, upper panel**). This is specifically complemented by suppression of factors involved in PTEX-complex, Mauer’s Clefts and membrane trafficking and protein export to the host cell cytoplasm but also several pathways of amino acid biosynthesis during *maturation* of the 5dMDPs (**Figure 2E**). The model of the specific metabolic and physiological state is also supported by the gene set enrichment analysis (GSEA) of the steady state transcriptome of the d5MDPs reflecting the enrichment of cellular and metabolic pathways along that of mRNA levels (**Figure 3B**). Essentially genes involved in formation of exosomes, centrosome, DNA repair machinery and nucleosome assembly exhibit the highest levels of expression in 5dMDP, along with factors of mitochondrial DNA replication and cytochrome oxidase biogenesis and a specific subset of exported proteins. Broadly, these transcriptional profiles were highly reproducible on both the *maturation* and *persistence* phase of the 5dMDPs, respectively (**Figure 3C and Figure S5 and S6**). Overall, these findings support our original hypothesis that *P. falciparum* dormancy is facilitated by highly specific transcriptional profile reprogramming, “supplying” the parasite with a unique set of biological functions. This re-programming mediates all three phases of dormancy including its *maturation, persistence* and ultimately *re-emergence* to asexual growth.

### The d5MDPs are more resistant to antimalarial compounds compared to the IDC stages

Taken together, it is feasible to suggest that the d5MDPs represent the true dormant stages of *P. falciparum* that give rise to recrudescent malaria infection after treatment with artemisinins and possibly other conditions. As such there are several main considerations that are important to understand regarding the d5MDPs viability and sensitivity to (other) antimalaria drugs. To address these, first we studied d5MDPs viability by investigating the intactness of mitochondrial membrane potential (MMP). Using the fluorescence microscopy-based detection of MMP, *P. falciparum* was stained with the MitoTracker™ Deep Red dye (**Figure 3D**). Curiously, essentially all pyknotic parasite cells at both day 1 and day 5 post DHA exposure stained with MitoTracker™. A quantitative analysis of the MitoTracker™ staining intensity indicated that on both day 1 and day 5, parasites exhibit somewhat lower fluorescence intensity compared to ring stage parasites which exhibited a wide range of mitochondrial staining (**Figure 3D, boxplot**). This may reflect the fact that even though the dormant parasites possess an enlarged mitochondrion and induce expression of mitochondrial genes, the MMP is somewhat compromised as a result of oxidative damage (see discussion).

Being viable even on day 5 after induction, it is now feasible to investigate the level and profile of sensitivity of the d5MDPs to antimalarial drugs. To do so, we carried out a drug sensitivity assay in which d5MDPs were matured for four days and subsequently treated with a series of antimalarial drugs or inhibitory compounds at concentrations equivalent to their 1x, 10x and 100x of the 50% inhibitory concentrations (IC_50_) determined for IDC asexual parasite stages (^IDC^IC_50_, **Figure S7**). After 24 hr exposure (completed on day 5), the drug was removed and re-initiation of the asexual growth was monitored for up to 30 days (**Figure 4**). Altogether, 1x ^IDC^IC_50_ treatments of d5MDPs with most tested antimalarial drugs did not alter their recovery rate suggesting their full resistance to these concentrations. The exceptions were amodiaquine (AMQ) and trichostatin (TSA) that caused a delay in recovery by 5.6 days and 4.8 days on an average respectively at 1x ^IDC^IC_50_. Furthermore, d5MDPs appear resistant to even 10x and 100x ^IDC^IC_50_ concentrations of atovaquone (ATQ), pyrimethamine (PMT), triclosan (TRI). There was an intermediate effect of 10-100x ^IDC^IC_50_ of DHA, MMV 1576856, (spiroindolone class) and TSA delaying the asexual growth recovery by 5-10 days. Finally, d5MDPs appear sensitive to several classes of antimalarials including mefloquine (MFQ), lumefantrine (LMF) amodiaquine (AMQ) and doxycycline (DOX) at 10-100X ^IDC^IC_50_. In particular, 100x ^IDC^IC_50_ concentrations of MFQ, LMF and AMQ that still correspond to their physiological concentrations, blocked the asexual growth recovery of the d5MDPs completely (past the 24 day observation period) or delayed by more than 20 days (**Figure 4**). In addition to these, we tested the d5MDP toxicity of tafenoquine (TAF) which has high efficacy against the *P. vivax* liver dormant stage hypnozoites. Due to the need of *in vivo* metabolic conversion, the drug has a high *in vitro* ^IDC^IC_50_ that reach their physiological concentration (TAF ~500nM, **Figure S7**). At this concentrations, tafenoquine didn’t affect d5MDP reemergence. Although 10x and 100x ^IDC^IC_50_ of TF did block the growth recovery to some extent, these concentrations are clinically unrealistic, and the observed *in vitro* effect likely represents (some form of) general toxicity. It has been previously observed, that adding gibberellic acid (GA) directly to the DHA-treated parasites promotes the faster recrudescence (M. Duvalsaint *et al*., 2018). Here, we used three different concentrations of GA (1 μM, 10 μM and 100 μM) on d5MDPs. However in our case, GA didn’t affect the recrudescence and its delay of d5MDPs (**Figure 4**).

**Fig 4.**
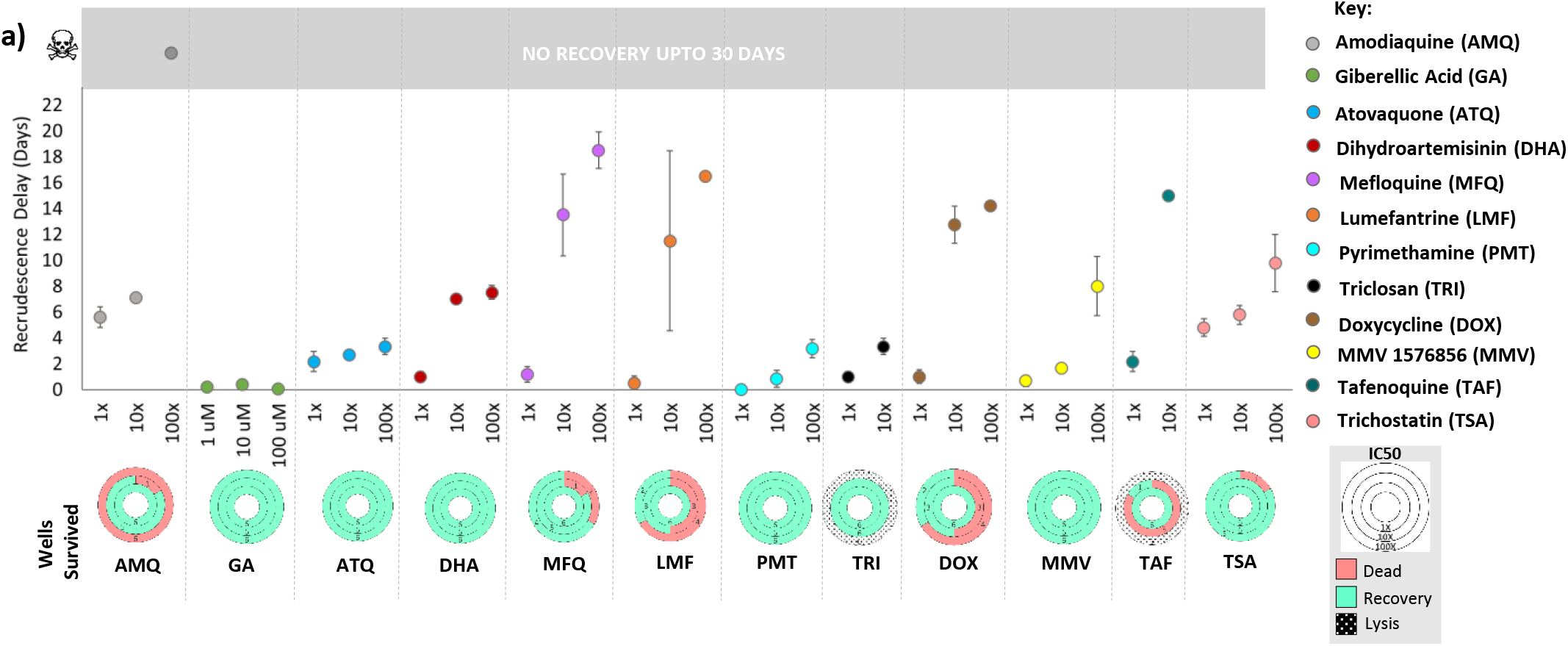
Drug sensitivity profiling of *P. falciparum* d5MDPs. A graph showing recrudescence delay for d5MDPs treated with various antimalarial drugs/compounds for 24-hours (started on Day 4) as compared to the controls (i.e. parasites treated with the corresponding drug solvent – DMSO, methanol or water, see **Figure S7**). Three different concentrations (i.e. 1xIC_50_, 10xIC_50_, 100xIC50) were tested per drug. Three independent biological replicates were conducted with two technical replicates per condition. Error bars indicate SEM calculated from all the cultures/wells that eventually recovered. The circular charts below show the number of cultures/wells which recovered, died (i.e. no recovery up to 30 days) or lysed for each drug concentration tested (legend on the right).

### Cellular structure characterizations of d5MDPs as a new *P. falciparum* life cycle developmental stage by transmission electron microscopy

Next, we studied the ultrastructure of d5MDPs by performing transmission electron microscopy post DHA treatment on day 1, day 5 (d5MDPs) and asexual ring stages for comparison. As shown in **Figure 5A and Figure S8**, the ring stage parasites show cuplike shape with well-defined sub-cellular organelles such as the nucleus, rough endoplasmic reticulum (RER), ribosome and membranous organelles such as mitochondria (M) and Golgi apparatus (GA). In contrast, 24hrs after DHA exposure (Day 1), the ring stage parasites transform into round shaped forms with somewhat lucent areas in the cytoplasm harboring fewer ribosomes and the presence of organelles such as the GA, RER and M still identifiable (**Figure 5B and Figure S8)**. d5MDPs, however, exhibit a highly irregular morphology characterized by a lack of intracellular organellar structures, presence of large “empty” vacuole-like structures (VS) and swollen mitochondria. There is also reduction in the number of ribosomes in the cytoplasm, but on the other hand, massive formation of multi-membrane structures (MAS) (**Figure 5C**). Several of these features have been reported previously in quinine, artesunate and/or piperaquine treated *P. falciparum*^31,32^, however most of these ultrastructural studies described short-term (hours) morphological changes upon drug exposure in the trophozoite stage. It is also important to note that we also observed frequent electron dense formations in the infected RBCs cytosol possibly indicating “packets” of proteins exported by the parasite (**Figure 5D and Figure S9**). Strikingly, these “packets” were observed mainly in infected RBCs in day 5 samples (**Figure S9 and S10)** and not uninfected RBCs, possibly indicating an active reduction in protein processing, folding and export to host cell surface during dormancy. Overall, our results describe the long-term morphological changes in DHA induced pyknotic dormant stages of *P. falciparum* for the first time.

**Figure 5:**
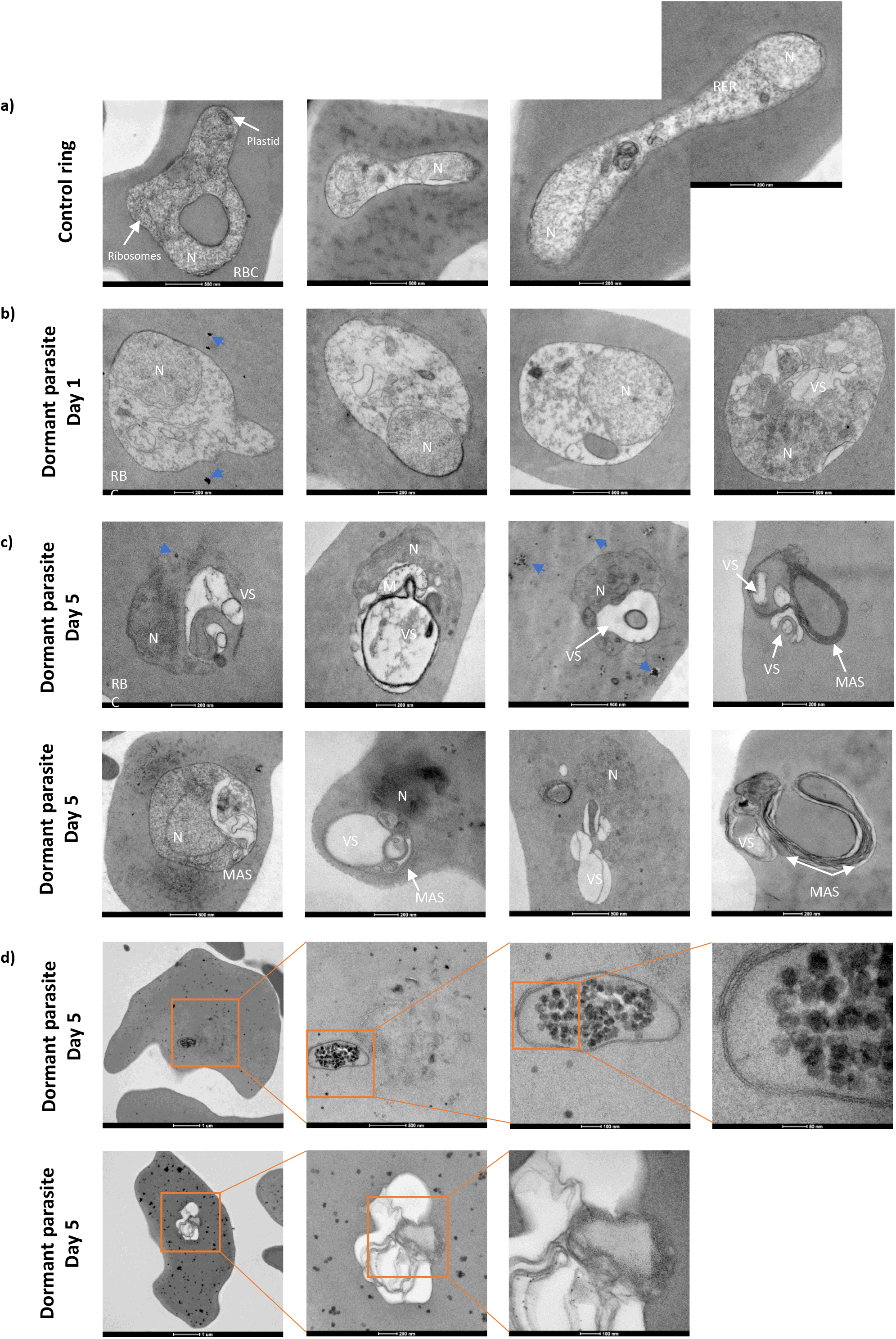
Transmission electron micrographs showing ultrastructures of *P. falciparum* ring stages and post DHA treatment on Day 1 and 5. **a)** Low-power transmission electron micrograph of ring stages with distinct organelles such as nucleus (N), ribosomes, plastid and rough endoplasmic reticulum (RER). **b)** Low-power transmission electron micrograph of dormant parasites 1 day post DHA treatment, showing the transformation of parasites to round and compact forms, with the nucleus (N) being the most identifiable organelle in the parasite. Note the presence of electron dense spots in the RBC cytosol (blue arrowhead). **c)** Low-power transmission electron micrograph of d5MDPs showing a completely different morphology characterized by lack of intracellular organellar structure except for the nucleus (N) and the presence of large “vacuole-like” structures (VS) and multi-membrane structures (MAS). An increase in electron dense spots (blue arrowhead) was also observed. **d)** Low-power transmission electron micrographs with increasing magnification (orange boxes) showing electron dense spots at Day 5 post DHA treatment were observed in the cytosol of 48 out of 65 infected RBCs (73%) on Day 5 (**Figure S10**). Additionally, clusters of these spots surrounded by membranous structures, forming “packets”, were observed in d5MDP infected-RBCs.

## Discussion

As opposed to the liver stage dormancy of *P. vivax* that has been a subject of numerous studies^33^, the existence of dormant parasite forms of *P. falciparum* induced upon drug exposure during the IDC is a relatively new concept that still raises many questions. However, given their potential to cause important clinical phenotypes such as recrudescence and drug resistance^17,34,35^, it is imperative to gain full understanding of their biological significance. As shown in the original study, exposures of ring stage parasites to clinically relevant concentrations of DHA (~700nM) can transform varying proportions of the parasite population into dormant stages with varying degree of recovery rates^19^. The commitment to dormancy and recovery depends largely on the timing and concentration of the initial drug exposure but also on the *P. falciparum* genetic background^36^. Consistent with this, here we used the 3D7 *P. falciparum* strain and demonstrated a highly reproducible pattern in which a vast majority of parasites transitioned to pyknosis after exposure of the mid ring stages (6-12 HPI) to 700 nM DHA for 6 hr (**Figure 1 & 2**). Subsequently, growth recovery occurred consistently at day 8-10 by which more than two thirds of the parasite population re-entered the IDC. Also, the dormant parasites induced in this study maintained the mitochondrial potential indicated by the MitoTracker™ staining. This is consistent with previous observations of the dramatically enlarged mitochondria with abnormal cellular arrangement^25^ and active transcription of at least 5 mitochondria-linked genes^24^. This suggests that our experimental system genuinely captures these previously postulated asexual dormant stages and provides a tangible model for their further characterizations.

The key observation made in this study is the unique transcriptional profile that portrays the dormant stages, likely reflecting their unique cellular physiology. Besides the mitochondria-linked functions, we observed transcriptional upregulations of genes involved in mitochondrial and apicoplast functions; biosynthesis of several amino acids; lipoic and shikimate acids; and nucleotide precursors; RNA biogenesis and histone modifications; all of which reflects an active metabolic state of the dormant stage parasites. This agrees and further expands on previous single gene-based observations showing several factors of lipid synthesis and pyruvate metabolism expressed in artemisinin induced dormant parasite^22^. On the other hand, there was a marked suppression of glycolysis as well as transcription (RNA Pol I & II-driven RNA synthesis); translation (protein synthesis, folding, trafficking and posttranslational processing); and factors involved in the host cell cytoplasm remodeling (such as the PTEX translocon and Maurer’s clefts) all of which is essential for growth and development of the ring stages along the IDC^37^. Altogether, these results are consistent with initial suggestions that the blood stage dormant parasites suppress active growth but retains some degree of metabolic activity^25^. Canonically, this is equivalent to the generic state of growth arrest in eukaryotic cells collectively known as cellular senescence. Originally discovered in 1960’s^38^, cellular senescence is associated with growth arrests while maintaining significant metabolic activities^39^. This is typically accompanied by broad gene expression changes facilitating the senescence state^40^ (and resistance to apoptosis^41^). In higher eukaryotes, cellular senescence is triggered by shortening of telomere, during aging^42^. However, senescence can be also induced by external stresses, typically involving DNA damage mediated by reactive oxygen species (ROS) produced by abnormal activity of the mitochondria^43^. Here, a feedback loop between the DNA damage and abnormally functioning mitochondria can maintain senescence via elevated levels of ROS indefinitely^44^. In addition to the abnormally acting mitochondria (see above), here we observed high transcriptional levels of genes involved not only in DNA repair machinery but also chromatin structure such as histone chaperones and factors of nucleosome assembly (**Figure 2B & 3B**). During the IDC, these functions are typically upregulated in the schizont stage along with the factors of DNA replication; presumably as a quality control mechanism during DNA synthesis^37^. In the induced dormant stage, however, the high expression of the DNA repair and chromatin remodeling is uncoupled from DNA replication whose expression is suppressed. From these results it is tempting to speculate that analogously to higher eukaryotic cells, in *P. falciparum*, the artemisinin induced senescence-like state is maintained by an interplay between abnormally functioning mitochondria and DNA damage. On the other hand, in mammalian cells this feedback loop is mediated by a CDK inhibitor genes CDKN1A (p21)^44^. While this gene is not conserved in the *P. falciparum* genome (author’s note), several cyclin-dependent kinase genes were shown to be upregulated in the dormant parasites^23^. In the future studies it will be interesting to identify a possible putative regulatory factor that has the potential to mediate cellular senescence in *Plasmodium* parasites. It will be particularly interesting to investigate whether the induction of blood stage dormancy share common features with the preerythrocytic stages (hypnozoites) but also with the dormancy of other eukaryotic species including plants^45^.

Curiously, both mitochondrial genes and DNA repair factor were also implicated in artemisinin MOA and/or resistance^46–49^. Currently there are two opposing studies implicating (or not) dormancy in artemisinin resistance directly. *In vitro P. falciparum* parasite line resistant to artelinic acid exhibit higher sensitivity to dormancy but also higher recovery rate^36^. On the other hand, presence of the artemisinin resistance-linked PfK13 mutations does not seem to modulate dormancy after artemisinin induction and/or sorbitol treatment^50^. Indeed, the role of the dormant blood stage *P. falciparum* parasite in the disease progression and treatment outcome is still a matter of debate^34,51^. In the recent opinion article, Wellems T *et al* (2020) argued that the persister parasites are unlikely to be associated with the increased parasite clearance halftimes (^1/2^PC), the current phenotypic feature of artemisinin resistance^34^. Indeed, there appears to be no association between ^1/2^PC and recrudescence and longer ^1/2^PC do not cause delayed resolution of malaria infections^52^. On the other hand, there is mounting evidence that the artemisinins dormant parasites do occur *in vivo;* in humans^27^ as well as in rodent models^26^. In the latter study, growth recovery time was directly correlated to the dose of dormant parasite transferred to the rodent host suggesting their significance in recrudescence. Hence even though, dormancy might not be a direct factor in artemisinin resistance mediated by PfK13 mutations, it may play significant role in the currently emerging and increasingly spreading frequency of ACT failures. Here we demonstrated that after their maturation, the d5MDPs exhibit resilience to a broad spectrum of antimalaria drugs. Hence, 4-5 days after the ACT administration, when the short-lived artemisinin component of an ACT is eliminated from the blood, a patient could carry hidden reservoirs of dormant parasites with significantly decreased drug sensitivities. At this stage, it is the long-lasting partner drug of the ACT (e.g. MFQ, AMQ of LMF) that is expected to eliminate the remaining parasites. Consequently, it is feasible to speculate that in cases where the putative reservoirs of dormant parasites are not effectively eliminated by an ACT, the parasites have the chance to recrudesce. From that end, we propose that future ACT combinations take into consideration partner drugs with higher capacity to eliminate the 5dMDPs to prevent recrudescence along with increasing frequencies of ACT failures. One of such strategy includes the design of triple artemisinin combination therapies (TACT) that is currently being evaluated for its ability to prevent the rise of resistant parasites^53^.

Interestingly, genes with the highest expression levels in the d5MDPs *per se* encode factors of extracellular signaling including the *components of exosomes* and (*infected*) *erythrocytes derived microvesicles*. These were complemented by high expression of a specific subset of the *P. falciparum* exportome and factors of vesicular trafficking. Biological significance of *P. falciparum* derived exomes and microvesices, also collectively termed the infected red blood cell-derived external vesicles (iREV), for malaria parasite pathophysiology was only recently recognized^54^. The iREV were shown to exert their effect on several immune cell types by which the parasite can modulate the host immune response and as such its virulence and disease outcome (reviewed in^55–57^). Moreover, the iREV can be also readily endocytosed by parasitized erythrocytes which suggests their role in cell-to-cell communications within the parasite population^58^. Such communication allows to sense parasite density and thus regulate a delicate balance between virulence (asexual growth) and transmissibility (commitment to gametocytogenesis). The high expression levels of iREV-related factors observed in our study suggests that the dormant parasites may be highly active in such regulation. In the future studies it will be interesting to explore the role of the dormant parasite-derived iREVs in extracellular signaling along with the molecular mechanism regulating and facilitating the iREV secretion.

The formation of pyknotic Rings and dormant parasites during the drug treatment was associated with a drastic modification of parasite morphology. The parasite cell structure was completely modified with a round shape easily identified by optical microscopy (Giemsa-stained thin film)^59^ and the formation of vacuole-like structure and larger mitochondria after 24 hr of DHA exposure as described previously^25^. The large vacuole-like structure could be associated to a reshape of the digestive vacuole and associated cytostomal vesicles^60^. Interestingly, we identified the formation of electron dense spots mainly at day 5 post DHA treatment. Clusters of these spots surrounded by membranous structures, forming “packets” could be associated with the reshaping of parasite plasma membrane (PPM) and parasitophorous vacuolar membrane (PVM). The downregulation of *Plasmodium* translocon for exported proteins (PTEX), HSP70 chaperone cycle^61^ PV membrane (PVM) proteins^62^, exported proteins in Maurer’s clefts and early-transcribed membrane proteins (ETRAMPs) genes were linked to proteins associated with the PPM and the PVM. Three-dimensional transmission electron microscopy (3D-TEM) will be performed in the near future to fully understand the formation of *Plasmodium falciparum* dormant rings.

## Methods

### Parasite culture and Dormancy induction

3D7 strain (MR4) of *Plasmodium falciparum* was cultured in RPMI 1640 medium (Gibco) containing purified human packed RBCs, supplemented with 0.25% Albumax II (Gibco), 0.1 mM hypoxanthine (Sigma), Sodium bicarbonate (Sigma) (2 g/L), and gentamicin (Gibco) (50 μg/L). Cultures were maintained on the shaker in 37°C with 5% CO_2_, 3% O_2_, and 92% N_2_ in 2% hematocrit. Culture medium was replenished regularly at least every 24 hours, fresh RBCs were added once a week. Blood smears fixed with 100% Methanol (Fisher Chemical) and stained with 5% Giemsa (Sigma) for 20 minutes were used for microscopy examination. For parasitemia calculation, 500 RBCs were counted in at least five view fields and if parasitemia was below 1%, ~10 000 RBCs were counted instead.

Parasite culture containing ring stages was synchronized in two consecutive cycles prior to the start of the experiments via 5% sorbitol (Sigma) to achieve maximum age difference ~6h.

Dormancy was induced by adding 700nM Dihydroartemisinin (DHA) to the 6-12 HPI parasite rings for 6h. After this incubation period, the RBCs were washed 3x with warm RPMI media to remove the DHA thoroughly. Mature parasites unaffected by the treatment were removed by passing the culture through magnetic column (MACS) for three consecutive days after the drug treatment, as described elsewhere^19^.

### Flow cytometry, sorting strategy and RNA sequencing

Staining of the samples for the flow cytometry was performed as described elsewhere^28^. Briefly, 1μL (for well-plate) or 10μL (for tube) of *P. falciparum-infected* RBCs were added to the 50μL (well-plate) or 200μL of PBS respectively, containing 5μg/mL Dihydroethidium (DHE) (Sigma) and 8μM Hoechst 33342 (Thermo Scientific). DHE was used as a redox indicator and Hoechst for DNA visualization. Samples were incubated for 30 minutes in 37°C. Additional 200μL (well-plates) or 3mL of PBS were added after the incubation.

Fortessa X20 (BD) was used to acquire all flow cytometry data using the UV laser (355 nm excitation) for Hoechst 33342 detection and the blue laser (488 nm excitation) for the DHE. 50 000 events were recorded for each sample and FlowJo (Tree Star) was used for FACS data processing. Uninfected RBCs were stained as a negative control.

500 infected RBCs were sorted into 3μL of Lysis buffer (final concentration of RNAseOut 2U/μL and 1mg/mL BSA, Molecular Biology Grade (NEB), diluted into sterile PBS) by FACSAria II (BD) into 0.2mL PCR strip tubes. Sorted samples were spun briefly and processed afterwards or stored at −80 °C. To confirm the correct sorting and gating strategy, 3 000 - 5 000 cells were collected from each gate into 3μL of RPMI media. 1μL of packed blood was diluted into 9μL of RPMI media in the separate tube to prepare ~1 million RBCs/1μL. Then, 1μL of diluted blood was added to the tube with the sorted cells to achieve~0.3-0.5% parasitemia. The tube was briefly spinned afterwards and a short blood smear was made and stained by Giemsa to evaluate the stage of the sorted parasites in the corresponding sample (**Figure S1C**).

cDNA libraries were prepared by SMART-Seq2 protocol as stated elsewhere^30^, with certain adaptations for the dormant stages of *Plasmodium*. Sequencing libraries were generated by Illumina Nextera XT kit and DNA/RNA UD Indexes (Illumina) following the manufacturer’s protocol, the successful tagmentation and library size was verified by Bioanalyzer High Sensitivity DNA chips (Agilent). Libraries were pooled and sequenced on Illumina NovaSeq 6000 platform, generating ~8Gb data output per sample.

### *In vitro* Drug susceptibility assays

Three biological replicates of 6-12 HPI *Plasmodium* culture at 1% parasitemia, 2% haematocrit were exposed for 48 hours to 11 different concentrations of selected drugs (**Figure S7**) in two-fold serial dilution manner in 96-well plate format. Each biological replicate was performed in two technical replicates. 0.1% DMSO (Sigma) was used as a negative control. After the 48h drug treatment, cultures were washed 3x with fresh warm RPMI media. The growth of the parasites was determined after 72 hours by Flow Cytometry with Hoechst 33342 and DHE staining as stated above. For all the compounds, IC_50_ values of each individual biological replicate were determined by nonlinear regression analysis in GraphPad Prism 8. These IC_50_ values of all biological replicates for a specific drug treatment were then averaged and IC_50_ range interval ± SD was defined (**Figure S7**).

### Double drug recovery assay

6-12 HPI rings were treated by 700nM DHA for 6 hours on Day 0. Removal of mature IDC stages by MACS was conducted on Days 1-3. Culture was then treated on Day 4 with second drug for 24 hours (**Figure S7**) with three different concentrations, 1xIC_50_, 10xIC_50_ and 100xIC_50_. Drugs were washed three times by fresh warm RPMI on Day 5, followed by the addition of fresh blood. Fresh blood was subsequently added every 7 days to prevent blood lysis. 1μL of culture was stained with Hoechst 33342 and DHE for flow cytometry on Days 8-12 and everythree following days until Day 30 (50 000 events recorded). Recovery day of each specific well in well plate was determined as a higher percentage of trophs and schizonts in culture than a background value in uninfected blood controls determined by FACS. If the positive recovery reading was followed by a negative one, the culture wasn’t considered as recovered (false positive). Recovery delay was calculated as a subtraction of recovery day and negative control recovery day (culture treated with 700nM DHA on Day 0 for 6 hours and with drug solvents added for 24 hours on Day 4 instead of a drug itself → DMSO, H2O, MetOH). Cultures without any trophs or schizonts on Day 30 were considered as not recovered.

### Live cell imaging

10μL of infected RBCs were washed with PBS, followed by staining with 100nM Mitotracker Deep Red (Thermo Scientific; abs/em ~644/665nm) and 8μM Hoechst 33342 (Thermo Scientific; abs/em 350/460-490nm) for exactly 30 minutes. Cells were washed 3x with PBS afterwards, diluted with PBS into ~10% haematocrit and pipetted on the glass slide, covered with cover slip and fixed by a nail polish. Images were captured with Zeiss LSM980 confocal microscope equipped with Airyscan 2 detector using Plan-Apochromat 100x/1.46 oil objective. Images were processed using Zen 3.4 software (Carl Zeiss).

### Data analysis and clustering

Trim Galore program was used to remove amplification primers, adapters and low-quality bases from 3’ ends from obtained raw sequencing reads. Alignment to the *P. falciparum* genome was performed by HISAT2. Properly oriented paired reads mapped to the unique location of the genome were considered for counting. BEDTools software was used to calculate gene-specific read counts. Fragments per kilobase per million mapped reads (FPKMs) were used for further transcriptomic analysis. Fourier analysis on each expression profile of the transcripts was performed as described previously^37^. In brief, using fft() function in R, the Fourier phase was deducted for each gene’s expression profiles and sorted in ascending order to achieve the phaseogram. Clustering of the genes was performed by ClusterW program. TreeView was used for the heatmaps conception.

### Electron Microscopy

15μL of packed RBCs from *P. falciparum* cultures (post DHA treatment on day 1, day 5 and control ring stages) were fixed overnight in 2.5% glutaraldehyde (Ted Pella, Inc.) in PBS, pH 7.3, washed in buffer, post-fixed for 1hr in 1% osmium tetroxide with 1.5% Potassium Ferrocyanide in PBS, block stained for 1hr in aqueous 1% uranyl acetate, then dehydrated in an ethanol series, absolute acetone and embedded in Araldite 502 resin (Ted Pella, Inc). Ultra-sections were cut at 90nm on a Leica EM UC6 Ultramicrotome, collected on 200 mesh copper grids covered with a Formvar carbon support film (Electron Microscopy Sciences), and stained for 8 mins in lead citrate stain. Photographs were taken with Transmission Electron Microscopy (Tecnai Spirit G2, FEI, USA) at 100kV with a bottom mounted digital camera FEI Eagle (4k x 4k pixels).

## Supplementary Figures

### Supplementary Figures Legends

**Figure S1.**
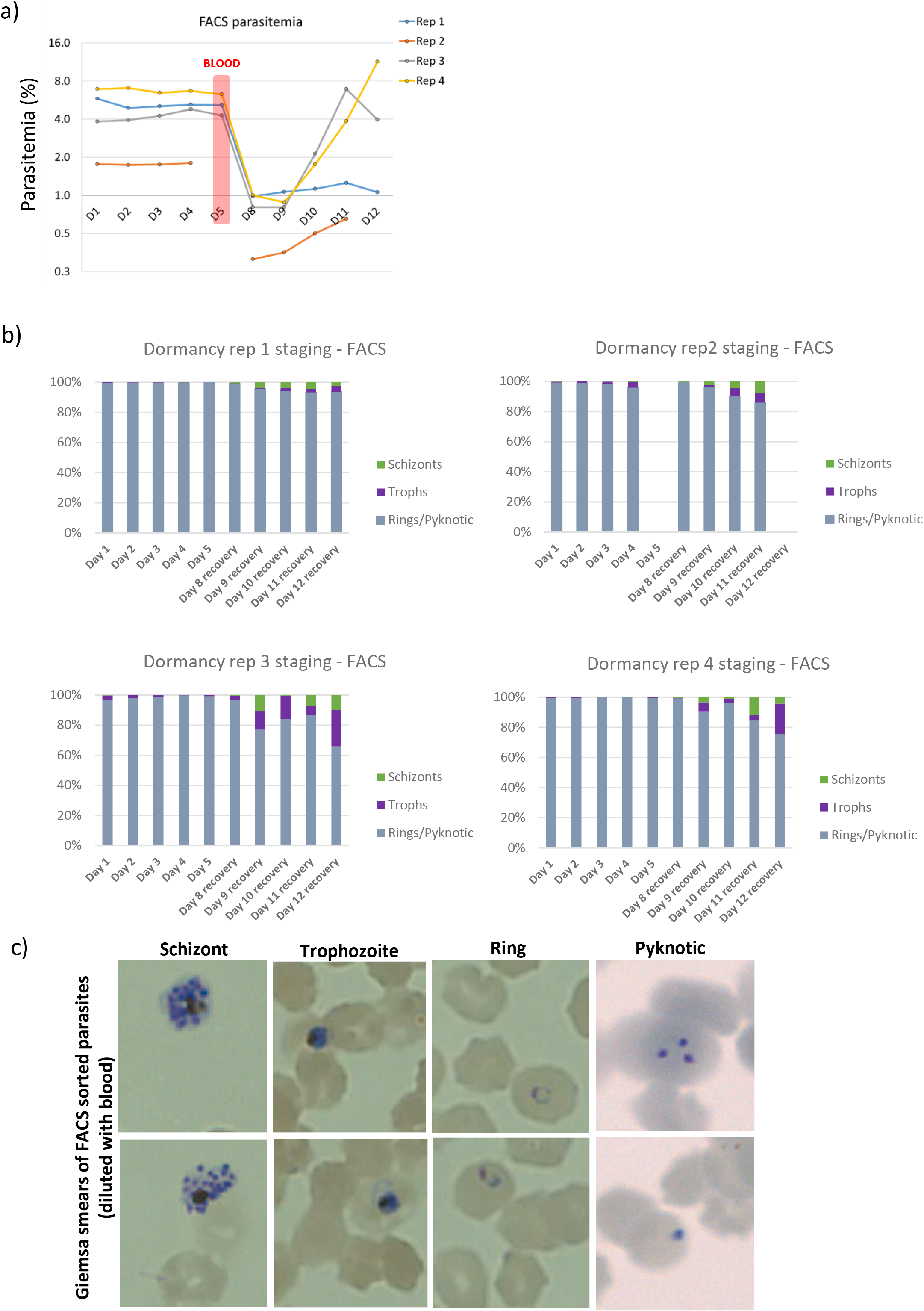
Parasitemia and staging based on flow cytometry after 700nM DHA exposure. **a)** Parasitemia (%) following the exposure of *P. falciparum* ring stages upon treatment with 700nM DHA (*induction*) and *re-emergence* of the dormant parasites up to Day 12. 50,000 events were collected using FACS to determine the parasitemia values. Blood was added on Day 5 to prevent the lysis of RBCs. Day 5 values could not be recorded for one of the replicates (Rep 2). **b)** Proportion of different parasite stages evaluated by FACS in individual biological replicates. 50,000 events were collected every day to estimate the stage proportions in the culture (processed using FlowJo™ software). **c)** Giemsa-stained blood smears were prepared for FACS sorted parasites (diluted in blood) from corresponding gates.

**Figure S2.**
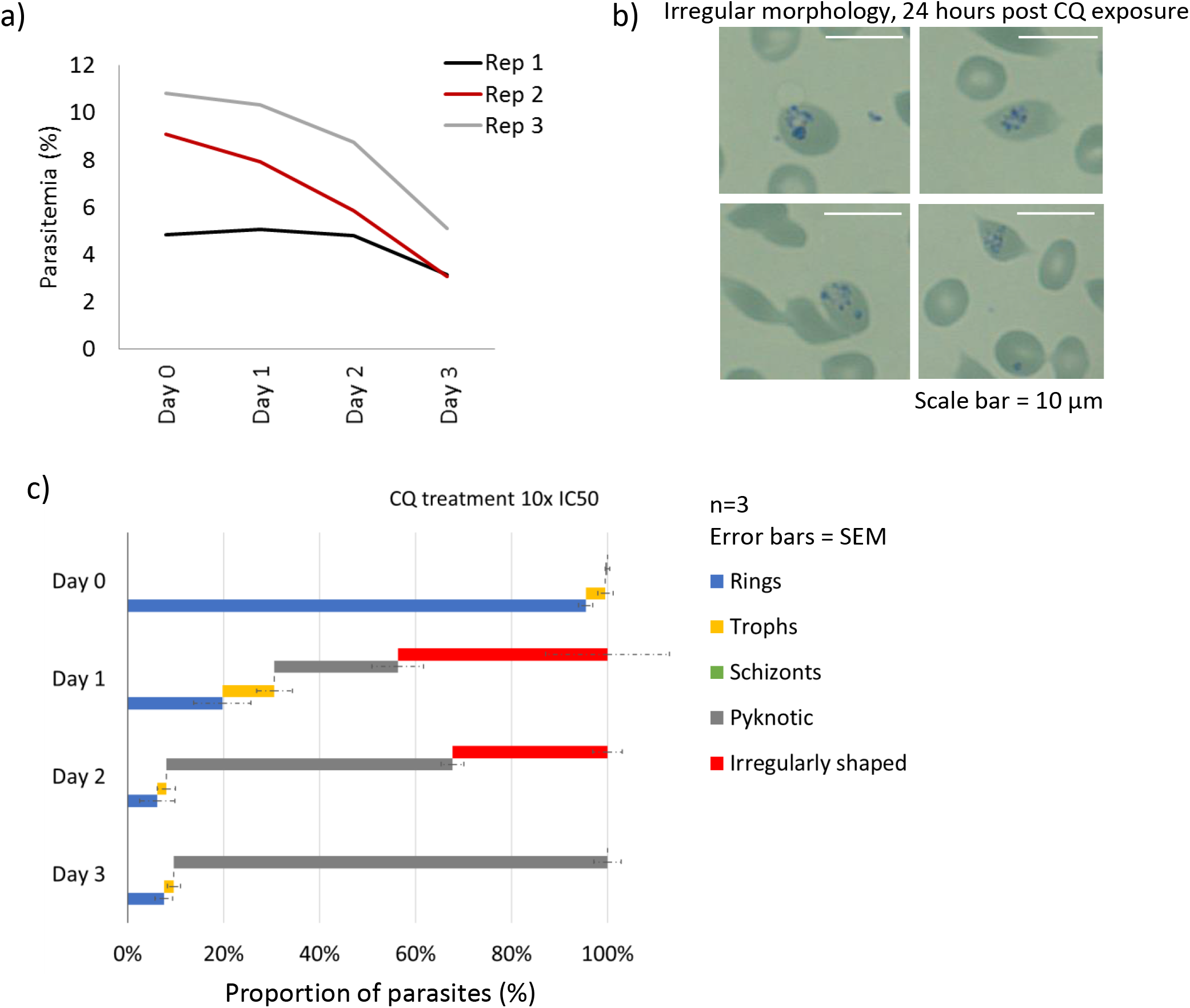
Parasitemia and parasite morphology upon 250nM (10xIC50) chloroquine treatment for 24 hours. **a)** Parasitemia (%) recorded on the first three days after chloroquine treatment of *P. falciparum* ring stage parasites in three independent biological replicates. Giemsa smears were used for the evaluation, where at least 500 RBCs were counted per condition. **b)** Example of the irregular morphology of the chloroquine-treated parasites after 24 hours, stained by Giemsa. Scale bar = 10 μm. **c)** Proportion of different parasite stages found in chloroquine-treated parasites evaluated by microscopy using Giemsa stained blood smears. 300 cells were counted for every day in three independent experiments. Error bars represent SEM.

**Figure S3.**
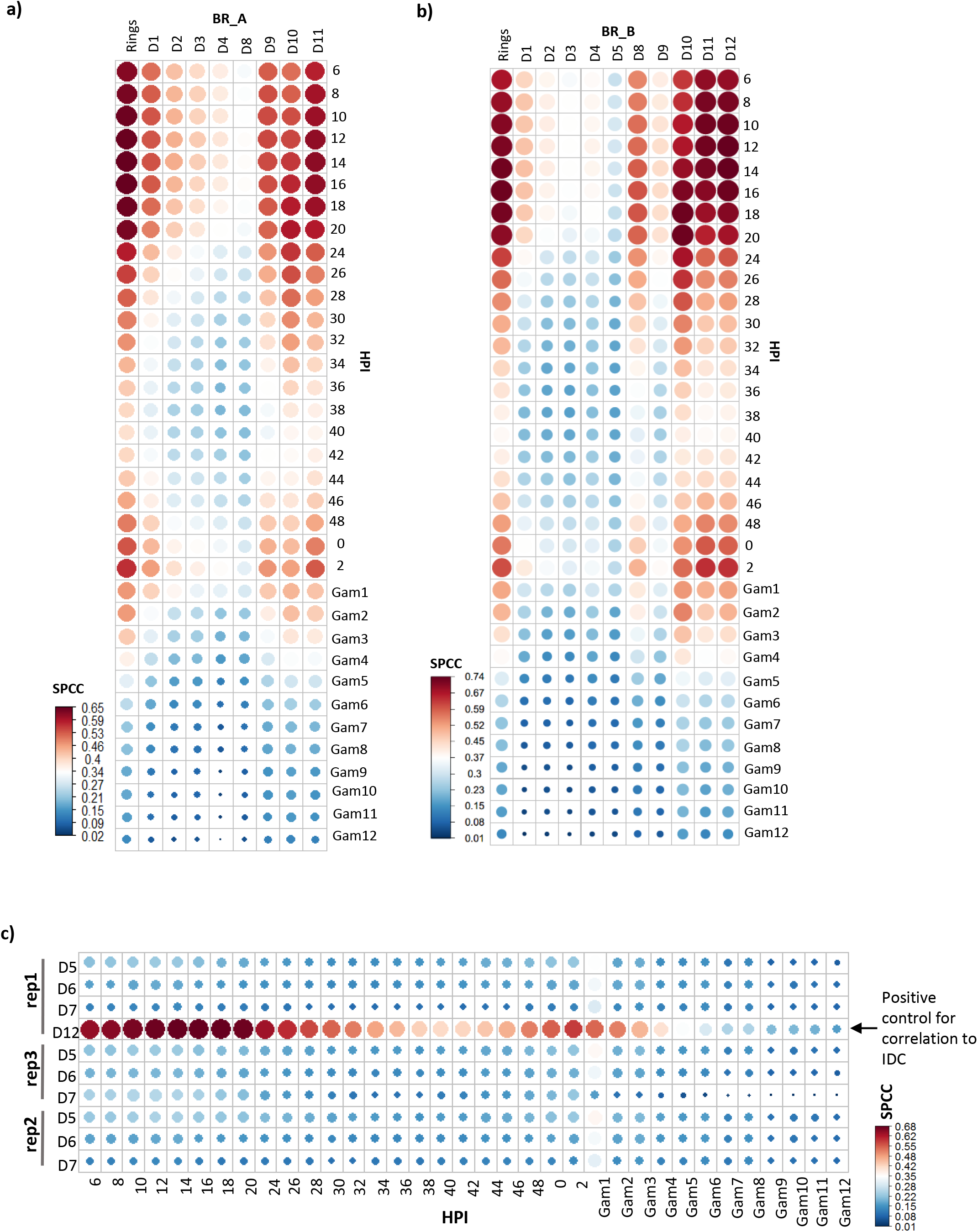
Correlation of Dormancy Developmental Cycle replicates to 3D7 *P. falciparum* reference transcriptome. Bubble plots depicting Spearman’s Correlation Coefficient (SPCC) for **a)** biological replicate A (“BR_A”), **b)** biological replicate B (“BR_B”), **c)** Day 5,6,7 *persistence* phase replicates, correlated to 0 to 48 HPI asexual IDC timepoints and gametocyte transcriptome. Red (larger circles) and blue (lower circles) colour depicts high and low correlation values, respectively.

**Figure S4.**
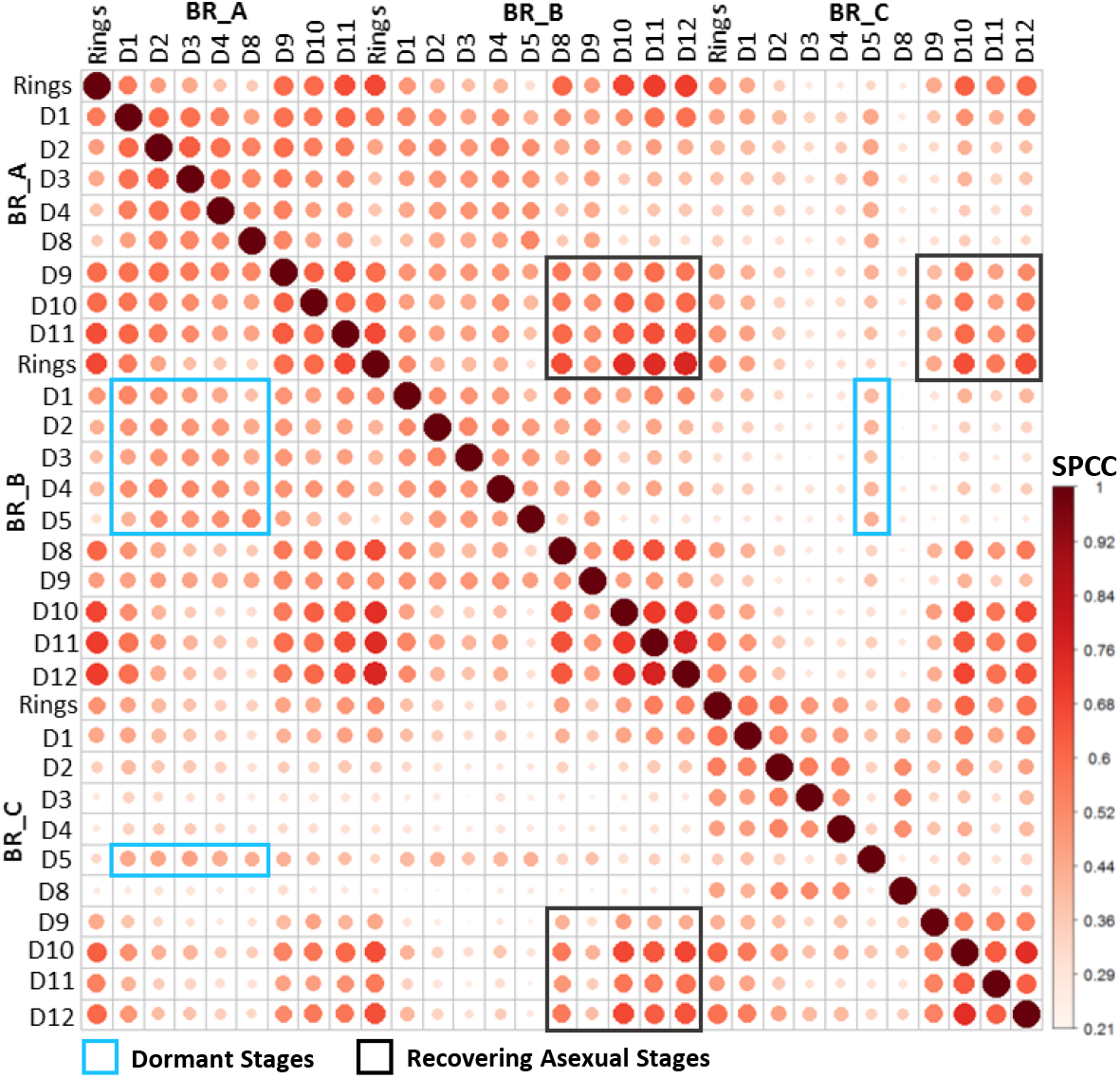
Correlation of all three biological replicates of the Dormancy Developmental Cycle transcriptome in DHA treated *P. falciparum*. A bubble plot depicting SPCC values for transcriptomes generated on each day in three independent biological replicates, correlated to each other.

**Figure S5.**
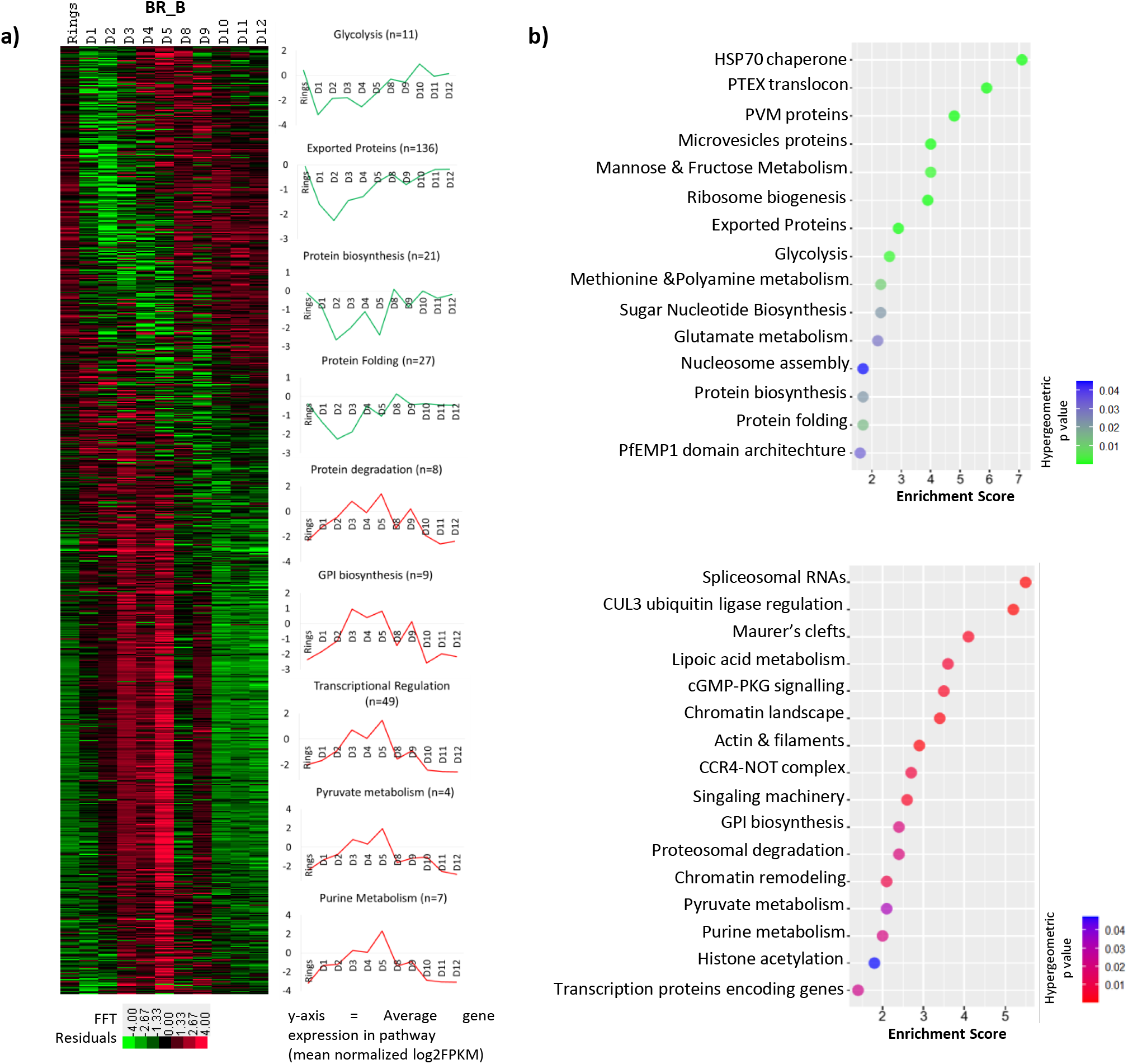
Transcriptomic profiling of dormancy developmental cycle in DHA treated *P. falciparum*. **a)** A heatmap depicting the Fourier transformed gene expression profile of *P. falciparum* during the *induction* and *re-emergence* phase of DHA induced dormancy in biological replicate B (“BR_B”). Genes upregulated and downregulated are highlighted by red and green respectively. Line graphs showing expression of key enriched pathways during Day 1 to 5 (downregulated in green, upregulated in red) are shown on the right panel. **b)** Bubble plots depicting pathways enriched amongst genes upregulated (lower panel) and downregulated (upper panel) during Day 1 to 5 post DHA induced pyknosis. All pathways are ordered by enrichment score and the color (red and green) denotes the statistical significance (hypergeometric p-value).

**Figure S6.**
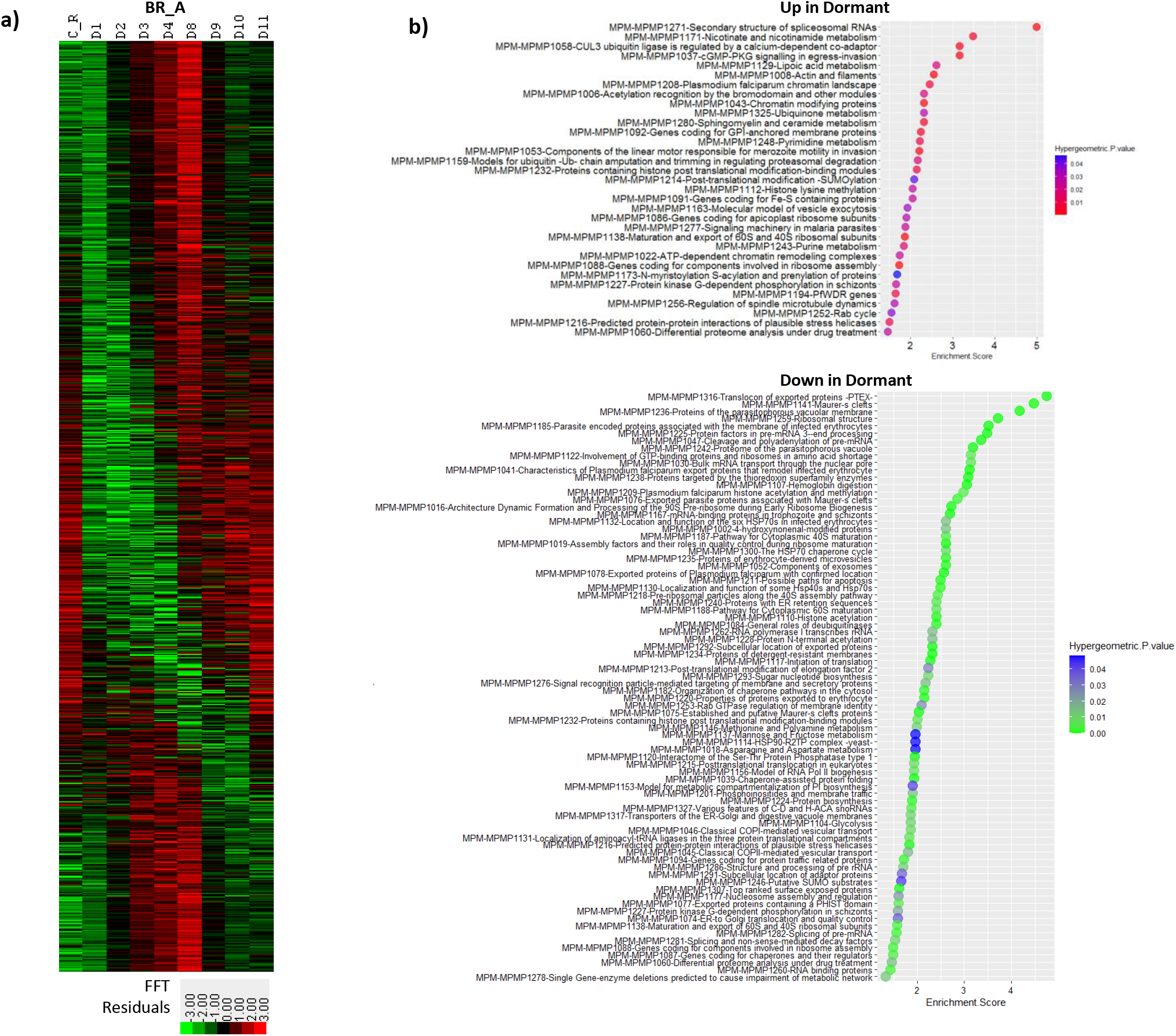
Transcriptomic profiling of dormancy developmental cycle in DHA treated *P. falciparum*. **a)** A heatmap depicting the Fourier transformed gene expression profile of *P. falciparum* during the *induction* and *re-emergence* phase of DHA induced dormancy in biological replicate A (“BR_A”). Genes upregulated and downregulated are highlighted by red and green respectively. **b)** Bubble plots depicting pathways enriched amongst genes upregulated (upper panel) and downregulated (lower panel) during Day 1 to 4 post DHA induced pyknosis. All pathways are ordered by enrichment score and the color (red and green) denotes the statistical significance (hypergeometric p-value).

**Figure S7.**
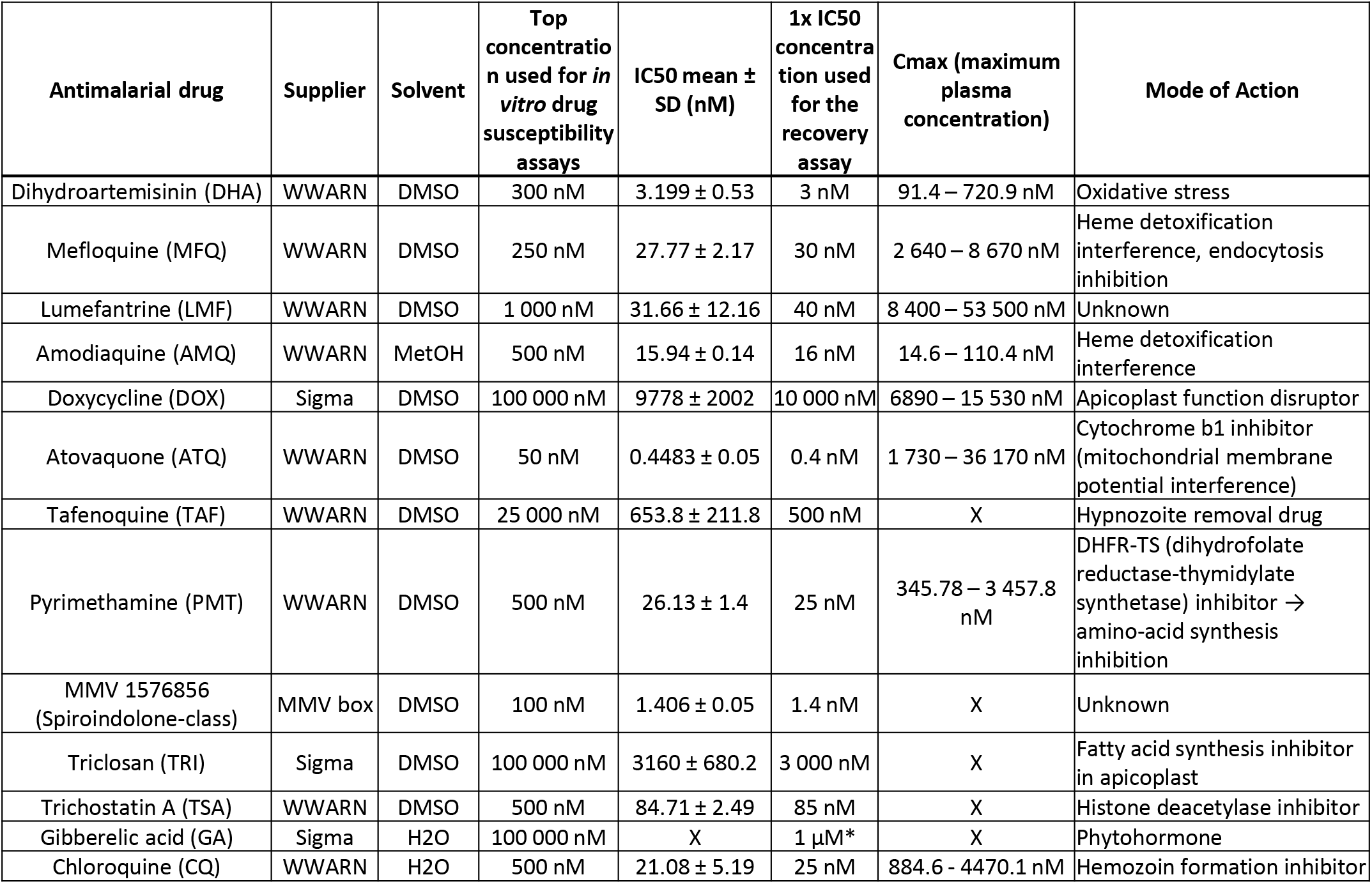
Antimalarial drug inhibitory concentrations and plasma concentrations. Table of compounds with IC50 concentrations used for the observation of changed recovery rate of d5MDPs (**Fig 4**). WHO-based maximum pharmacokinetic concentrations in human patients (reference – Guidelines for the Treatment of Malaria, 3^rd^ edition, WHO, 2015) are listed for comparison, together with the compounds’ mode of action.

**Figure S8:**
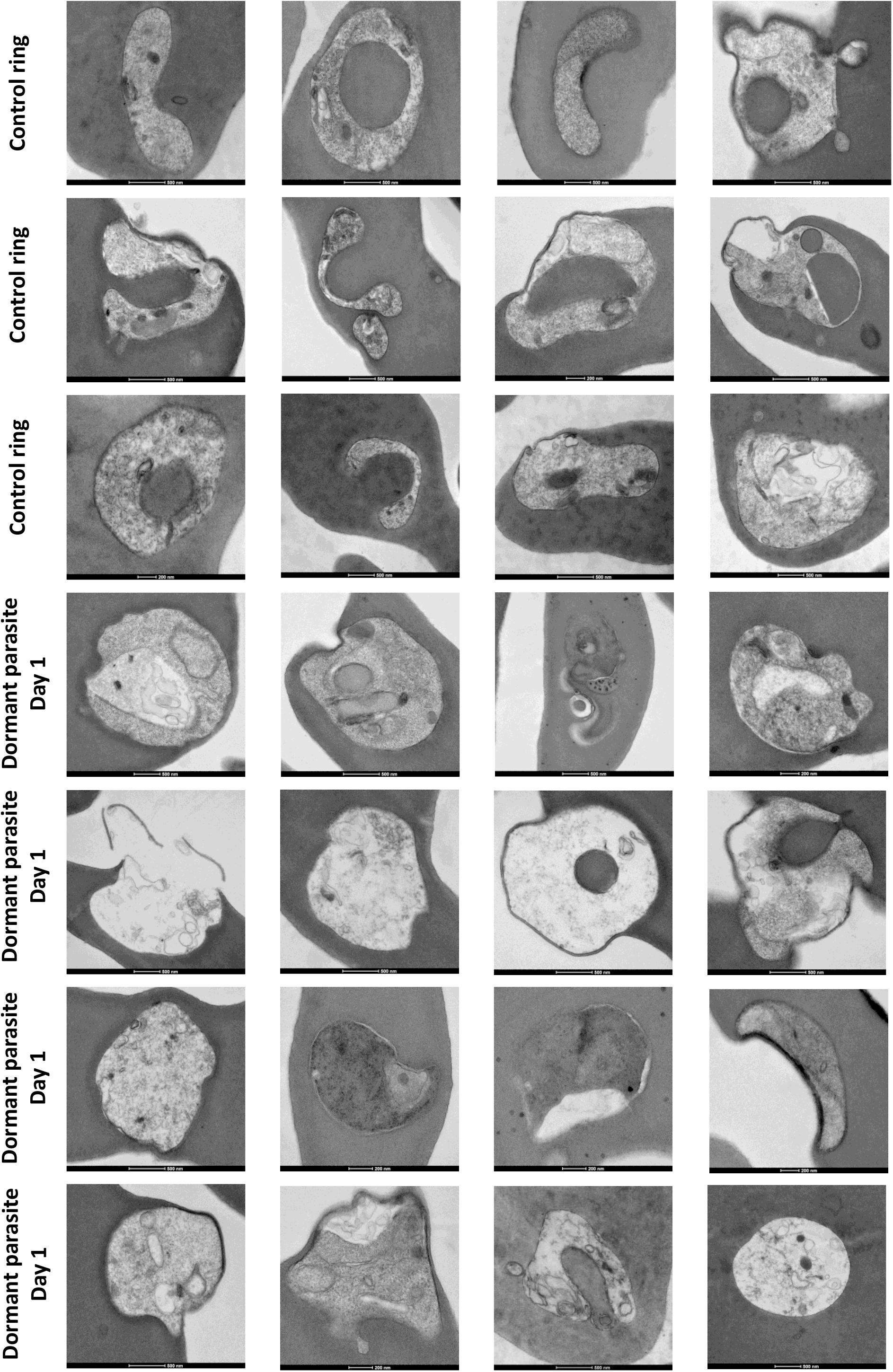
Additional TEM micrographs of untreated ring stages and DHA-treated parasites on Day 1. Transmission electron micrographs showing ultrastructures of *P. falciparum* ring stages and post DHA treatment on Day 1. Depending on the angle of microsections, the control rings were observed to be in a variety of forms: long oval, “C”-shaped and “O”-shaped. These ring forms transformed to round compact forms after 24 hrs post DHA treatment.

**Figure S9:**
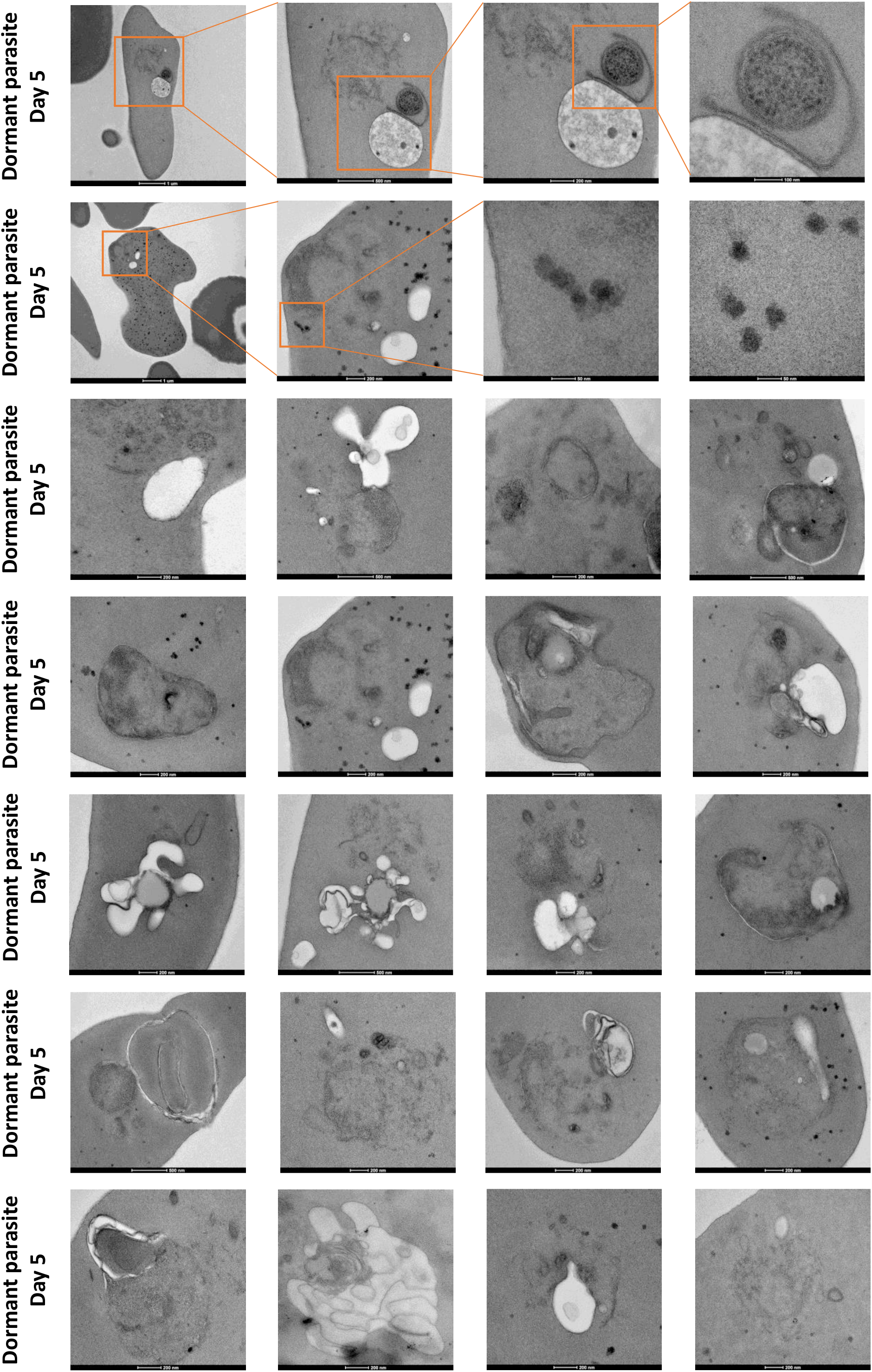
Additional TEM micrographs of d5MDPs showing electron-dense spots and packets. Transmission electron micrographs showing electron dense spots and “packets” in the cytosol of d5MDP-infected RBCs. Electron dense spots were observed either scattered through the cytosol or within membranous structures, here called “packets”, in the infected RBCs. Micrographs with increasing magnification (orange boxes) shows the close up details of these electron dense spots.

**Figure S10:**
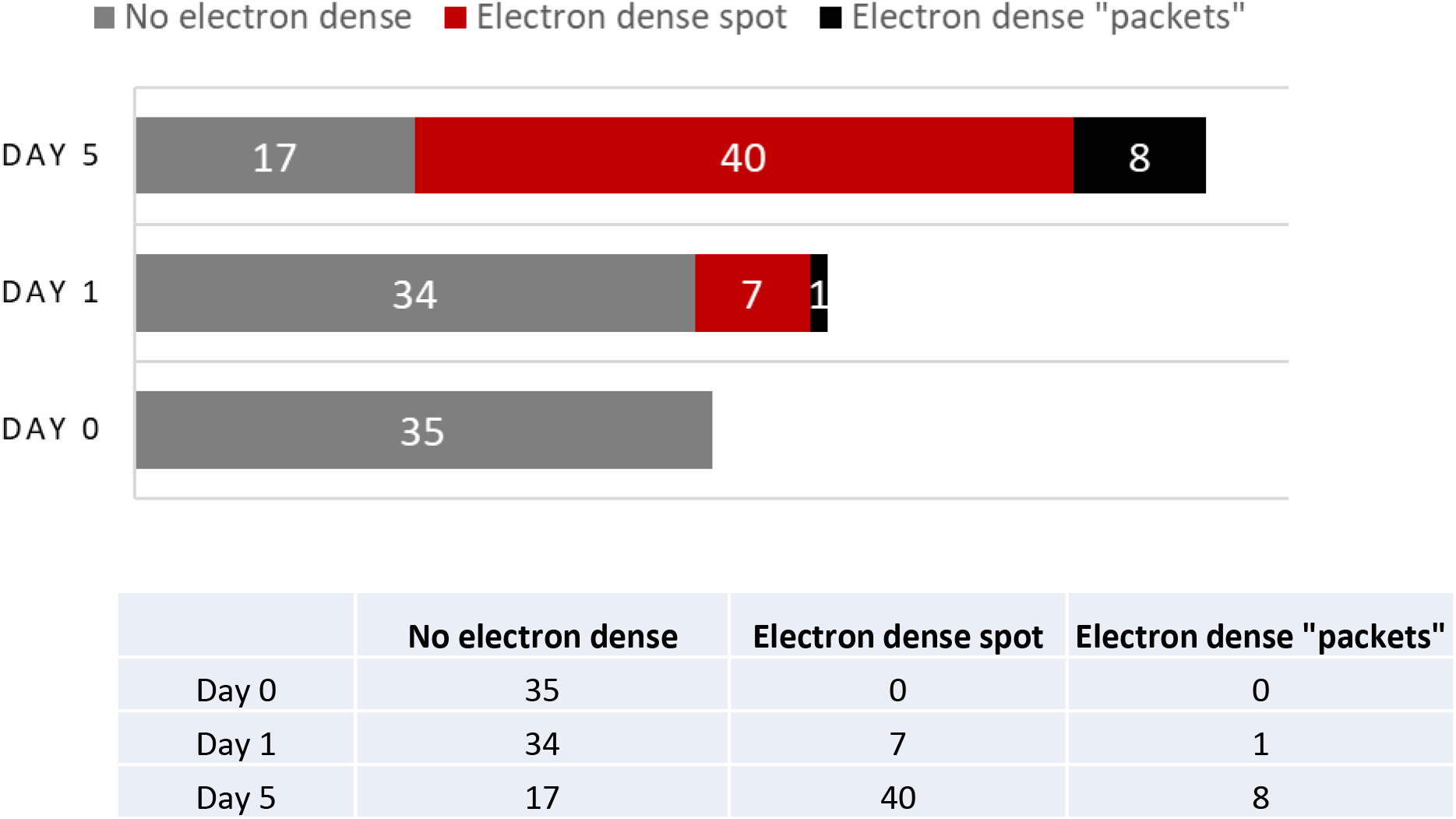
Quantification of Electron dense spots and “packets” in infected host RBC. A bar graph showing proportion of infected RBCs showing electron dense spots or “packets” in rings (Day 0), Day 1 and Day 5 post DHA treatment (d5MDPs). A table summarizing the same is shown below the graph.

## Notes

### Competing Interest Statement

The authors have declared no competing interest.

